# Choline Kinase Alpha Inhibition is a Mutation Agnostic Therapeutic Vulnerability in Clonal Hematopoiesis

**DOI:** 10.64898/2026.10.26.760729

**Authors:** Brandon J. Gheller, Olivia Mitchell, Michael Quach, Jayakrishnan Gopakumar, Serine Avagyan, Song Yang, Thomas P. Mathews, Chuck Haggerty, Alexander Meissner, Jacqueline S. Garcia, Ralph J. DeBerardinis, Siddhartha Jaiswal, Fernando Camargo, Leonard I. Zon

## Abstract

Clonal hematopoiesis (CH) affects the aging population and is associated with increased risk of hematological malignancy, cardiovascular disease, and all-cause mortality. Therapeutic options for CH remain limited. Using hematopoietic stem and progenitor cell (HSPC) color barcoding and mosaic mutagenesis in zebrafish, we identified perturbed choline metabolism and increased phosphatidylcholine (PC) biosynthesis via choline kinase alpha (CHKA) in *asxl1*-mutant HSPCs. Suppressing CHKA-dependent choline flux reduced *ASXL1*-driven clonal expansion. This effect was conserved in a murine *DNMT3A* and human *TET2* model. Mechanistically, CHKA-dependent choline flux controls phosphatidylcholine production that supports platelet activating factor (PAF) production. Exogenous PAF overcomes the suppressive effect of CHKA inhibition in *ASXL1* and *TET2* models. We isolated primary, human HSPCs from the peripheral blood of patients with endogenous *ASXL1* mutations and demonstrated that perturbing PAF signaling reduces their competitive potential in an *ex vivo* clonal competition assay. This CHKA/PAF axis is conserved in *TET2*-mutant HSPCs. Together, these findings identify a mutation agnostic metabolic dependency across the three most common CH genotypes and demonstrate that targeting the CHKA-dependent choline flux suppresses clonal expansion.

## Introduction

Human aging is accompanied by the accumulation of somatic mutations across tissues including hematopoietic stem and progenitor cells (HSPCs)^1^. When these mutations occur in hematological malignancy-associated genes they can confer a competitive advantage to individual HSPCs, resulting in clonal expansion a condition termed clonal hematopoiesis (CH)^2,3^. Approximately 6% of individuals 60-69 years old and 18.5% of those greater than 90 years of age have CH^2^. Over 75% of CH is driven by mutations in 3 genes: *ASXL1*, *TET2*, and *DNMT3A*^4^. CH increases the risk of hematological malignancy^5^, cardiovascular disease^6^, chronic liver disease^7^, chronic obstructive pulmonary disease^8^, and acute kidney injury^9^. Despite extensive investigation into CH biology, no therapies are currently approved to prevent or treat mutant clone expansion.

Emerging evidence demonstrates that metabolic abnormalities enhance the competitiveness of CH-driving clones. Metabolic profiling of human HSPCs with the hotspot DNMT3A mutation *DNMT3A*^R882^ or its murine correlate *Dnmt3a*^878H^ identified perturbed central carbon metabolism skewing these cells to a more oxidative phosphorylation phenotype^10–12^. Pharmacological inhibition of mitochondrial complex I by metformin was sufficient to quell the competitive advantage of *Dnmt3a*^878H/+^ cells *in vivo* by reversing these mutant clones’ metabolic and epigenetic abnormalities. These results were replicated using the mitochondrial-targeted antioxidant MitoQ^10^. Similarly, high doses of vitamin C can partially restore the function of wildtype TET2 in haploinsufficiency models^13,14^. There has been limited work identifying metabolic vulnerabilities in *ASXL1*-mutant clones. An unmet need remains to take an unbiased approach to metabolically profile CH-driving HSPCs and assess whether potential therapeutics can be identified that span genotypes^15,16^.

We had previously developed a system that combined an endogenous color barcoding system that labels unique HSPC clones (Zebrabow) with mosaic mutagenesis of CH-driving genes. This system was named <u>T</u>issue editing <u>W</u>ith Inducible <u>S</u>tem cell <u>T</u>agging via <u>R</u>ecombination (TWISTR)^17^. TWISTR allows for the competition of mutant and wildtype (WT) HSPCs without the need for confounding transplantation and allows for the prospective isolation of live clones. Using TWISTR, we identified phosphatidylcholine (PC) metabolism as a metabolic vulnerability in *asxl1*-driven CH. These findings were confirmed through mutagenesis of *asxl1* in primary, human HSPCs combined with d9-choline stable isotope tracing. Inhibiting PC metabolism by targeting choline kinase alpha (CHKA), the enzyme responsible for the phosphorylation of choline, reduced *asxl1*-mutation driven CH progression *in vivo* in zebrafish and *ex vivo* in primary, human HSPCs. Targeting CHKA was also shown to be effective in reducing the clonal expansion of *Dnmt3a*-mutant cells in a murine transplantation model and in a *TET2* primary, human HSPC *ex vivo* model. We developed a patient-derived HSPC clonal expansion assay to demonstrate that inhibiting CHKA was also effective in cells derived from patients with myelodysplastic syndromes (MDS). Mechanistically, we demonstrate that CHKA dependent platelet activating factor (PAF) supports HSPC proliferation of *ASXL1* and *TET2* mutant HSPCs.

### Clonal hematopoiesis is characterized by enhanced phosphatidylcholine biosynthesis

To evaluate the metabolic differences between clonally dominant HSPC clones and their WT counterparts we used the TWISTR system^17^. To employ the TWISTR system zebrafish embryos from crosses of *Tg(Zebrabow:M)* and *Tg(draculin:CreER^T^*^2^) were injected with a pool of validated gRNAs targeting CH related genes followed by induction of color barcoding with treatment of 4-hydroxytamoxifen at 24 hours post fertilization (hpf), the birth of the hematopoietic stem cell in zebrafish, and grown to adulthood (**Figure 1A**)^18^. We performed low-input, untargeted metabolomics on mutant, dominant HSPCs as defined by *>*40% of cells being derived from a single-color clone and compared this to HSPCs from a zebrafish injected with a gRNA known to not affect hematopoietic clonality. Of 90 identified metabolites, 7 differed significantly between dominant and non-dominant HSPC clones. Of these we filtered metabolites not previously identified in mammalian cells and this resulted in 30 metabolites, 4 of which were significantly different (**Figure 1B**). Notably, two choline-related metabolites were depleted in dominant clones – acetylcholine and betaine (**Figure 1B**, **Supplementary Figure 1A**)^19^. We then developed a targeted metabolite profiling assay to probe choline metabolism specifically allowing us to quantify 5 choline-related metabolites in a manner sensitive to distinguish between cells isolated from the HSPC and myeloid lineages in as few as 50 000 cells (**Supplementary Figure 1B**). In a cohort of zebrafish injected solely with a gRNA targeting exon 12 of *asxl1* (previously identified as the strongest driver of CH in zebrafish)^18^ our targeted choline metabolite profiling assay confirmed that betaine is significantly less abundant with acetylcholine following a similar pattern in dominant HSPC clones with choline being unchanged (**Figure 1C**). Therefore, we posited that choline metabolism was altered in dominant HSPCs to support their competitive potential.

**Figure 1.**
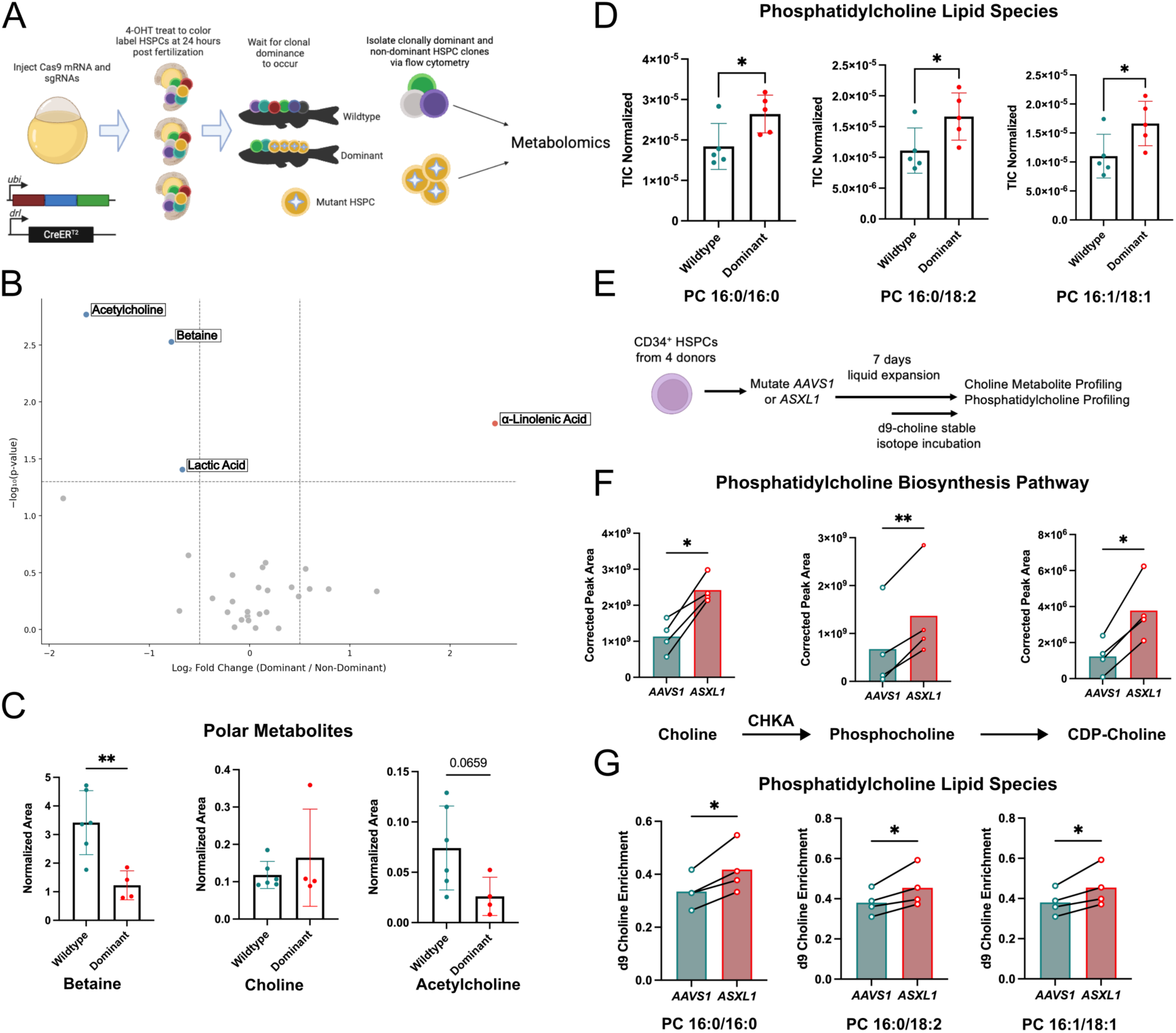
Hematopoietic Stem and Progenitor Cells (HSPCs) Harboring Clonal Hematopoiesis Driving Mutations are Characterized by Perturbed Choline Metabolism. A) Experimental schematic for metabolite profiling assay that combines color barcoding with mosaic mutagenesis to model CH. B) Volcano plot showing individual metabolite differences between dominant and wildtype (WT) HSPCs (n = 5 per group). C) Betaine, choline, and acetylcholine levels in HSPCs from an independent cohort of zebrafish with CH-driven by *asxl1* or WT control using a standard controlling metabolite profiling assay. **P < 0.01 (unpaired t-test). D) Quantification of three phosphatidylcholine species (16:0/16:0, 16:0/18:2, 16:1/18:1) in dominant (*asxl1*-mutant) and WT HSPCs in zebrafish *P < 0.05 (unpaired t-test). E) Experimental schematic for metabolite profiling and stable isotope tracing approach paired with *ASXL1* mutagenesis in primary, human HSPCs. F) Levels of polar metabolites in the phosphatidylcholine biosynthesis (choline, phosphocholine, and CDP-choline) pathway in primary, human HSPCs edited with CRISPR/Cas9 and a gRNA targeting a control region of the genome (*AAVS1*) or *ASXL1*, n = 4 biological replicates, *P < 0.05, **P < 0.01 (paired t-test). G) Levels of integration of d9-choline into phosphatidylcholine species 16:0/16:0, 16:0/18:2, 16:1/18:1 in primary, human HSPCs edited with CRISPR/Cas9 and a gRNA targeting a control region of the genome (*AAVS1*) or *ASXL1*, n = 4 biological replicates, *P < 0.05, **P < 0.01 (paired t-test).

To understand if the decrease in betaine was relevant to the development of clonal dominance, we developed an *in vivo* d9-choline stable isotope tracing approach paired with the TWISTR system (**Supplementary Figure 2A**). We noted that betaine biosynthesis from choline was increased in dominant HSPCs despite lower overall betaine levels (**Supplementary Figure 2B**). Betaine can serve as a methyl donor in certain tissues through the transfer of a methyl group to methionine via *BHMT* to form dimethylglycine and methionine (**Supplementary Figure 2B**)^20^. Previous studies have identified aberrant methylation patterns in HSPCs with CH-driving mutations such as *DNMT3A*^21^, and we also demonstrated, through whole genome bisulfite sequencing, that *asxl1*-mutant, dominant HSPCs have 2933 differentially methylated regions (**Supplementary Figure 3A**). We therefore hypothesized that the betaine to methionine conversion pathway may be activated in *ASXL1-*mutant HSPCs to support the observed altered methylation patterns. In d9-choline stable isotope tracing studies in primary, human HSPCs that had been mutated in *ASXL1* we noted no labelling from d9-choline in S-adenosylmethionine or methionine despite >50% of betaine being labelled (**Supplementary Figure 2C**). Altogether these findings demonstrate that although betaine levels were decreased, flux from choline to betaine was increased, isotope tracing excluded one-carbon metabolism as the major fate of this metabolic alteration.

Acetylcholine and betaine represent two metabolic fates of choline with the third being PC lipid species^19^ that would not be identified in our original polar metabolite profiling analysis (**Supplementary Figure 1A**). To this end, we performed a low input PC profiling approach on dominant and wildtype HSPCs in tandem with our TWISTR model and quantified 26 distinct PC species (**Supplementary Table 1**). *asxl1*-mutant, dominant HSPCs had a greater abundance of three specific PC species, 16:1/18:1, 16:0/18:2, and 16:0/16:0 (**Figure 1D**). We next evaluated if the increase in PC levels observed in *asxl1*-mutant, dominant HSPCs was conserved in humans. Following a published protocol^22^ that was shown to be effective in increasing HSPC expansion in liquid culture we mutagenized *ASXL1 (*or *AAVS1* as a control) in CD34^+^ HSPCs collected from peripheral blood of healthy humans after mobilization and expanded them for 7 days then repurified the CD34^+^ fraction for targeted metabolite profiling (**Figure 1E**). Choline as well as the two polar precursors to PC, phosphocholine and cytidine diphosphate-choline (CDP-choline), were all increased in *ASXL1-*mutant, human HSPCs (**Figure 1F**). Using an identical experimental design we added d9-choline 48 hours before CD34^+^ re-purification and performed targeted PC profiling to quantify active PC biosynthesis of the species we identified in the zebrafish. Mirroring our findings in zebrafish we identified that synthesis of PC(16:1/18:1), PC(16:0/18:2), and PC(16:0/16:0) were all increased in *ASXL1* mutant HSPCs (**Figure 1G**). Together these findings demonstrate a conservation of perturbed PC synthesis in HSPCs with CH-driving mutations across species.

### Choline kinase alpha inhibition suppresses hematopoietic clonal expansion

We hypothesized CHKA-dependent choline flux was required for clonal expansion of mutant HSPCs. To test this, zebrafish harboring similar size *asxl1* mutant clones as determined by variant allele frequency (VAF, **Supplementary Figure 4A**) were injected with an inhibitor of CHKA. CHKA phosphorylates choline to create phosphocholine in the first step of choline-PC synthesis^23^. RSM-932a (CHKAi) was chosen to inhibit CHKA-dependent choline flux as it is the most well described CHKA inhibitor^24^. CHKAi was injected intraperitoneally for 5 days and *asxl1* VAF was measured two months post treatment (**Figure 2A**). We noted no death when zebrafish received escalating doses of CHKAi from 0.1 μg/kg to 10 μg/kg. A 10 μg/kg dose of CHKAi had no adverse effects on hematopoietic output (**Supplementary Figure 5AB**) or number of stem cells contributing to hematopoiesis (**Supplementary Figure 5C**) and therefore, 10 μg/kg was used for all subsequent experiments. The VAF of the largest *asxl1* mutant clone was reduced 2 months post treatment in the peripheral blood (**Figure 2B**) demonstrating CHKAi abrogated progression of CH in zebrafish. Altogether these data identify CHKAi as a potential therapeutic option for CH with the need for cross-species validation.

**Figure 2.**
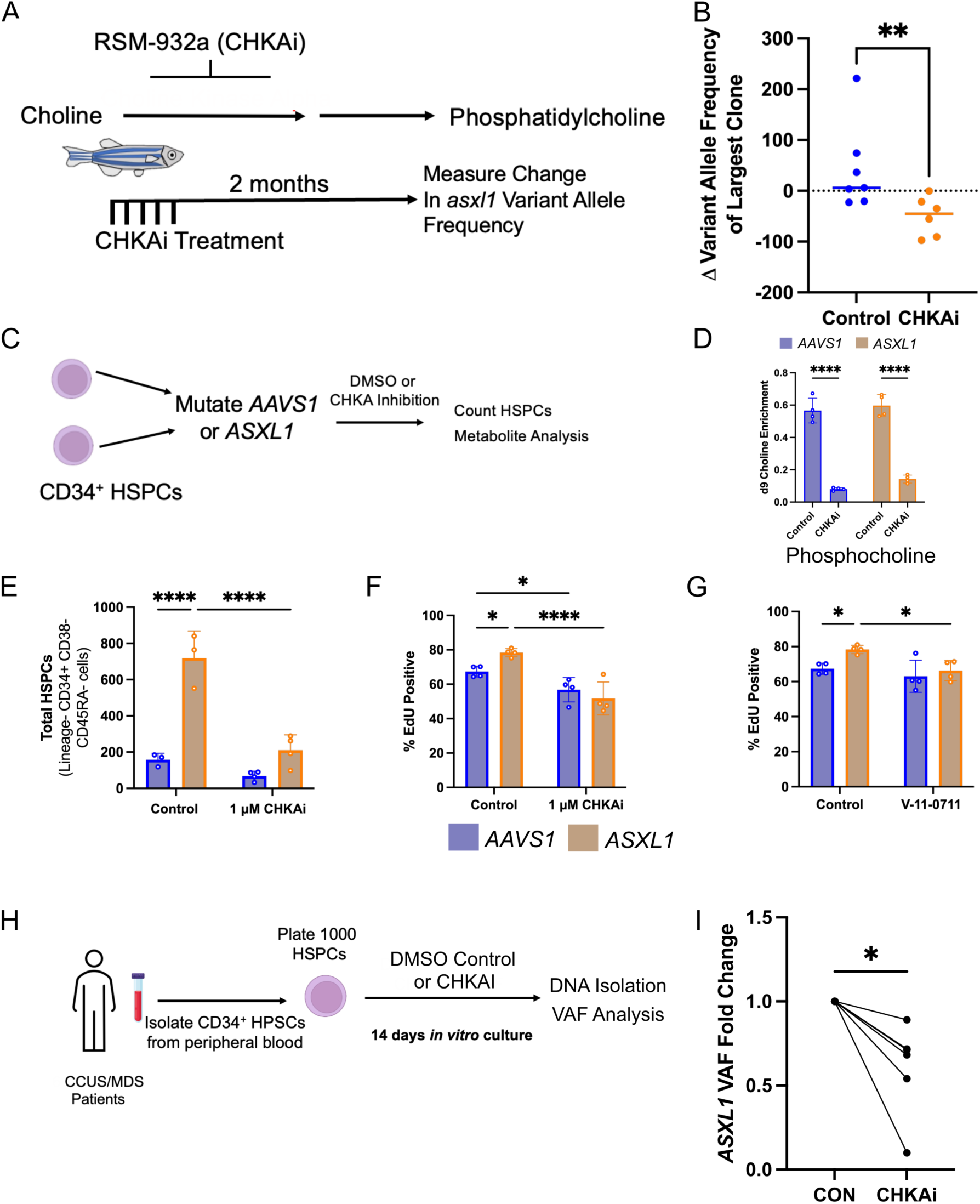
Choline Kinase Alpha Inhibition Suppresses *ASXL1*-mutant Clonal Outgrowth. A) Experimental schematic for treating zebrafish with RSM-932a (CHKAi). B) Quantification of the change of the largest *asxl1* variant allele frequency (VAF) in the peripheral blood 2 months after an initial treatment of PBS control or CHKAi, n = 6-7 biological replicates, **P < 0.01 (unpaired t-test). C) Experimental design for CRISPR/Cas9 mutagenesis of *AAVS1* or *ASXL1* primary, human HSPCs followed by CHKAi or control treatment and subsequent metabolic profiling or clonal expansion assays. D) Levels of integration of d9-choline into phosphocholine in primary, human HSPCs (CD34+ enriched) after 7 days of culture in drug treatment, n = 4 biological replicates, *P < 0.05, ***P < 0.001, ****P < 0.0001 (two-way repeated measures ANOVA followed by Tukey’s Post Hoc). E) Total number of primary, human HSPCs (Lin-CD34+CD38-CD45RA-) 14 days after sorting 1000 cells per well in response to 1 μM CHKAi, n = 3-4 technical replicates representative of 3 independent experiments, ****P < 0.0001 (two-way ANOVA followed by Tukey’s Post Hoc). F) EdU positivity of cultures of HSPCs 4 days after *AAVS1* or *ASXL1* mutagenesis, EdU was incubated for 24 hours from day 3 to day 4, in response to 1 μM CHKAi, n = 4 technical replicates representative of 3 independent experiments, *P < 0.05, ****P < 0.0001, (two-way ANOVA followed by LSD). G) EdU positivity of cultures of HSPCs 4 days after *AAVS1* or *ASXL1* mutagenesis, EdU was incubated for 24 hours from day 3 to day 4, in response to 1 μM V-11-0711, n = 4 technical replicates representative of 3 independent experiments, *P < 0.05 (two-way ANOVA followed by LSD). H) Experimental schematic of how human HSPCs were obtained from peripheral blood draws of patients with clonal cytopenia of undetermined significance (CCUS) or myelodysplastic syndrome (MDS) and clonal dynamics evaluated in response to CHKAi by variant allele frequency (VAF). I) Quantification of *ASXL1* VAF fold change between DMSO control and CHKAi 14 days after treatment initiation, n = 6 biological replicates, *P < 0.05 (paired t-test).

To model *ASXL1* mutation driven clonal expansion in human cells CD34^+^ HSPCs from healthy donors were obtained and mutated in *AAVS1*, as a control, or *ASXL1* and HSPCs (CD34^+^Lin^−^CD38^−^CD45RA^−^) were repurified and plated in a liquid media favoring HSPC expansion (**Figure 2C Supplementary Figure 6A**). Dose response studies showed that 1 μM of CHKAi was not toxic in human HSPCs (**Supplementary Figure 6B**) and that this dose did not influence differentiation based on colony formation output (**Supplementary Figure 6C**). The 1 μM dose was also effective in reducing d9-choline incorporation in the polar intermediates (phosphocholine and CDP-choline), in the biosynthesis of choline to PC species (**Figure 2D Supplementary Figure 6D**), and into PC species globally (**Supplementary Figure 6E**). *ASXL1* mutagenesis was sufficient to increase the number of HSPCs (CD34^+^Lin^−^CD38^−^CD45RA^−^) after 14 days of culture whereas CHKAi abrogated this effect (**Figure 2E**). EdU labeling at 3 days post mutagenesis showed that the increase in *ASXL1*-mutant HSPCs at 14 days was driven by an increase in proliferation at this early time point (**Supplementary Figure 6F**). CHKAi decreased the *ASXL1*-mutant cells’ proliferative advantage in the 3–4-day window demonstrating a decrease in *ASXL1*-mutation driven proliferation was the specific method by which CHKAi was reducing clonal expansion (**Figure 2F**). We used a second pharmacological agent, V-11-0711, to inhibit CHKA through an orthogonal mechanism^25^. This drug also reduced PC levels in human HSPCs (**Supplementary Figure 6G**) to validate the specificity of the effect of suppressing PC biosynthesis in reducing proliferation of *ASXL1*-mutant HSPCs. Similar to RSM-932a, V-11-0711 reduced EdU incorporation in *ASXL1*-mutant HSPCs (**Figure 2G**). In sum, pharmacological suppression of CHKA activity is sufficient to reduce proliferation of HSPCs harboring *ASXL1* mutations leading to a reduction in clonal expansion in zebrafish *in vivo* and in human cells *ex vivo*.

To evaluate the therapeutic potential of modulating CHKA-dependent choline flux we sought a system that faithfully recapitulated human, clonal disorders. To this end we developed an approach whereby CD34^+^ HSPCs are isolated from the peripheral blood of patients with somatic CH mutations who had not been exposed to mobilizing agents. These patients were identified as having either clonal cytopenia of undetermined significance or MDS (**Figure 2H Supplementary Figure 7A Supplementary Table 2**). This system allows for the quantification of clonal dynamics in response to treatment in mutant and non-mutant HSPCs from the same individual exposed to the same environment without the need for exogenous genetic manipulation. After 14 days of culture CHKAi treatment decreased *ASXL1* VAF indicating that more cells in the culture were WT thereby suppressing *ASXL1*-driven clonal expansion (**Figure 2I**). Importantly, there was no effect on total cell number in HSPCs derived from peripheral blood of a healthy volunteer obtained in a similar manner and exposed to CHKAi (**Supplementary Figure 7B**). The suppression of mutant HSPC clones *ex vivo* from patients suggest that CHKAi is able to operate in a manner that is not deleterious to WT clones allowing patients to still support normal hematopoiesis after treatment. Furthermore, these data suggest CHKAi may be valuable for hematopoietic clonal disorders beyond CH.

### CHKA inhibition is a conserved metabolic vulnerability in TET2 and DNMT3A mutant HSPCs

Because *ASXL1*, *TET2*, and *DNMT3A*, the three most common CH-driving mutations, alter distinct molecular processes we asked whether their shared competitive phenotype reflects a convergence on a common metabolic requirement. *TET2* was mutated using a previously validated gRNA and we identified this was sufficient to increase cell number after 14 days of culture and this enhanced clonal expansion was suppressed by CHKAi (**Figure 3A**)^22^. *TET2* mutant HSPCs also exhibited increased PC biosynthesis (**Supplementary Figure 8A**). Like *ASXL1*, *TET2* mutagenesis increased proliferation within 4 days post mutagenesis and both CHKA inhibitors tested, RSM-932a and V-11-0711, returned proliferation to levels similar to the *AAVS1* control (**Figure 3B**). Therefore, *ASXL1* and *TET2* mutant HSPCs share a similar targetable metabolic vulnerability manifesting in an increased dependence on CHKA-dependent choline flux, we next asked if this was also true of *DNMT3A*.

**Figure 3.**
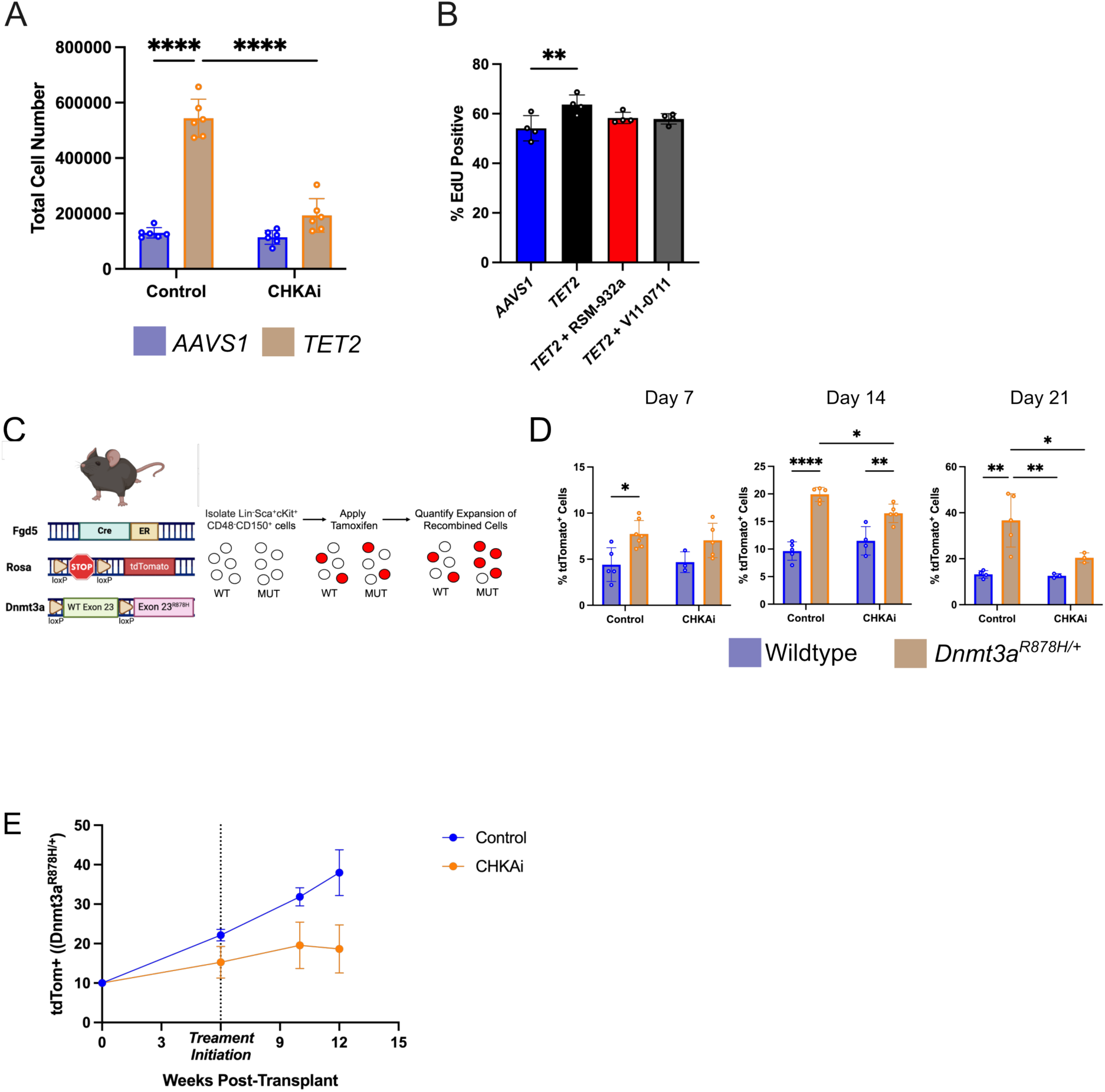
Choline Kinase Alpha Inhibition Suppresses *TET2* and *DNMT3A* Mutant Clonal Outgrowth. A) Total number of cells 14 days after sorting 1000 primary, human *AAVS1* or *TET2* – mutant HSPCs (Lin-CD34+CD38-CD45RA-) per well in response to 1 μM CHKAi, n = 6 technical replicates ****P < 0.0001 (two-way ANOVA followed by Tukey’s Post Hoc). B) EdU positivity of cultures of human, primary HSPCs 4 days after *AAVS1* or *TET2* mutagenesis, EdU was incubated for 24 hours from day 3 to day 4, in response to 1 μM CHKAi or V-11-0711, n = 4 technical replicates, **P < 0.01 (one-way ANOVA followed by Tukey’s Post Hoc). C) Experimental schematic for mosaically inducing mutant Dnmt3a^878H/+^ and tdTomato expression in isolated mice HSPCs (Lin-Sca-1+cKit+CD48-CD150+) *ex vivo.* D) Quantification of clonal expansion of *Dnmt3a^R^*^878^*^H/+^* mutant or wildtype control HSPCs in response to 1 μM CHKAi, n = 3-7 technical replicates, *P < 0.05, ** P< 0.01 (unpaired t-test). E) Percentage of cells in the peripheral blood of mice that are tdTomato/ Dnmt3a^878H/+^ 12 weeks post transplantation with or without weekly CHKAi injections, n = 3-7 biological replicates, *P < 0.05 (two-way ANOVA, main effect of treatment).

To test *Dnmt3a* driven clonal expansion, hematopoietic stem cells (Lin^−^Sca-1^+^cKit^+^CD48^−^CD150^+^) from mice harboring an inducible *Dnmt3a^R^*^878^*^H^* and tdTomato allele (*Dnmt3a*^878H/+^ Fgd5-CreER^T2^ LSL-tdTomato) or a wildtype control (*Dnmt3a*^+/+^ Fgd5-CreER^T2^ LSL-tdTomato) were isolated and cultured in a previously defined polyvinyl alcohol system (**Figure 3C**)^26–28^. Mosaic recombination and expression of tdTomato and *Dnmt3a^R^*^878^*^H^* was achieved through a moderate dose of tamoxifen resulting in <10% of cells being tdTomato^+^ in both the *Dnmt3a*^878H/+^ and *Dnmt3a*^+/+^ conditions at baseline. This results in a competitive system whereby tdTomato^+^/ *Dnmt3a*^878H^ compete against tdTomato^−^/ *Dnmt3a*^+/+^ in the same well and competition suppressive compounds can be determined based on suppressed expansion of the tdTomato^+^/ *Dnmt3a*^878H/+^ cells. *Dnmt3a^R^*^878^*^H^*^/+^ expression led to a clonal expansion of tdTomato^+^ cells which was suppressed by CHKAi at 14 and 21 days but expansion of tdTomato^+^ *Dnmt3a*^+/+^ cells treated with CHKAi was not different than control (**Figure 3D**). Similarly, CHKAi suppressed expansion of total cell number in the *Dnmt3a^R^*^878^*^H^*^/+^ group only (**Supplementary Figure 9A**). Mice were competitively transplanted with tdTomato^+^/*Dnmt3a^R^*^878^*^H^* cells. Six weeks after transplant mice were treated with 7.5 mg/kg CHKAi once weekly for 6 weeks based on a well-established protocol shown to be effective for reducing solid tumors in mice^24^. CHKAi significantly reduced the clonal outgrowth of tdTomato^+^/*Dnmt3a^R^*^878^*^H^*^/+^ cells in the peripheral blood (**Figure 3E**). Based on these results CHKAi presents a therapeutic option spanning the three most common CH-driving gene mutations.

### Enhanced CHKA-dependent choline flux is driven by a demand for platelet activating factor

To understand what might be underlying the demand of enhanced CHKA-dependent choline flux we performed RNA-seq on *asxl1* mutant, dominant HSPCs from zebrafish and prioritized enzymes known to be involved in downstream metabolism of PC. Interestingly, the only differentially expressed genes were downregulated and were generally involved in the recycling of lysophosphatidylcholine to PC (e.g., *enpp2, pld3, gpcpd1*)^29,30^ except for one upregulated gene, *lpcat2* (**Figure 4A, Supplementary Table 3**)^31^. LPCAT2 catalyzes the terminal acetylation step of the platelet activating factor (PAF) remodeling pathway converting lyso-PAF to PAF using a pre-existing PC pool. Separately, PAF can be synthesized *de novo* via direct condensation of CDP-choline with 1-O-alkyl-2-acetylglycerol which is a reaction that directly consumes the CHKA-dependent CDP-choline pool^32^. CDP-choline production is upregulated in *ASXL1*-mutant, human HSPCs and relies directly on CHKA-dependent choline flux and we determined its biosynthesis to be reduced by CHKAi (**Figure 1F and Supplementary Figure 6D**). Furthermore, CDP-choline also facilitates the secondary PAF *de novo* synthesis mechanism^32^. We next performed RNA-seq of human CD34^+^ HSPCs 7 days after *ASXL1* mutagenesis and while *LPCAT2* was the most upregulated *LPCAT* enzyme it was not significantly different between genotypes (**Supplementary Figure 10A**) although we did note an increase in chromatin accessibility in both identified promoter regions (**Supplementary Figure 10B**). *ASXL1*-mutant HSPCs did however have an increase in expression for the single PAF receptor *PTAFR* (**Figure 4B**) which was also accompanied by an increase in chromatin accessibility in both known promoter regions of the *PTAFR* gene (**Figure 4C**) which translated into increased protein expression (**Figure 4D**)^33^. Therefore, we hypothesized that *ASXL1* mutation driven increases in CHKA-dependent choline flux were facilitating enhanced clonal expansion through an increase in PAF synthesis and that concordant expression of PTAFR in mutant HSPCs was required.

**Figure 4.**
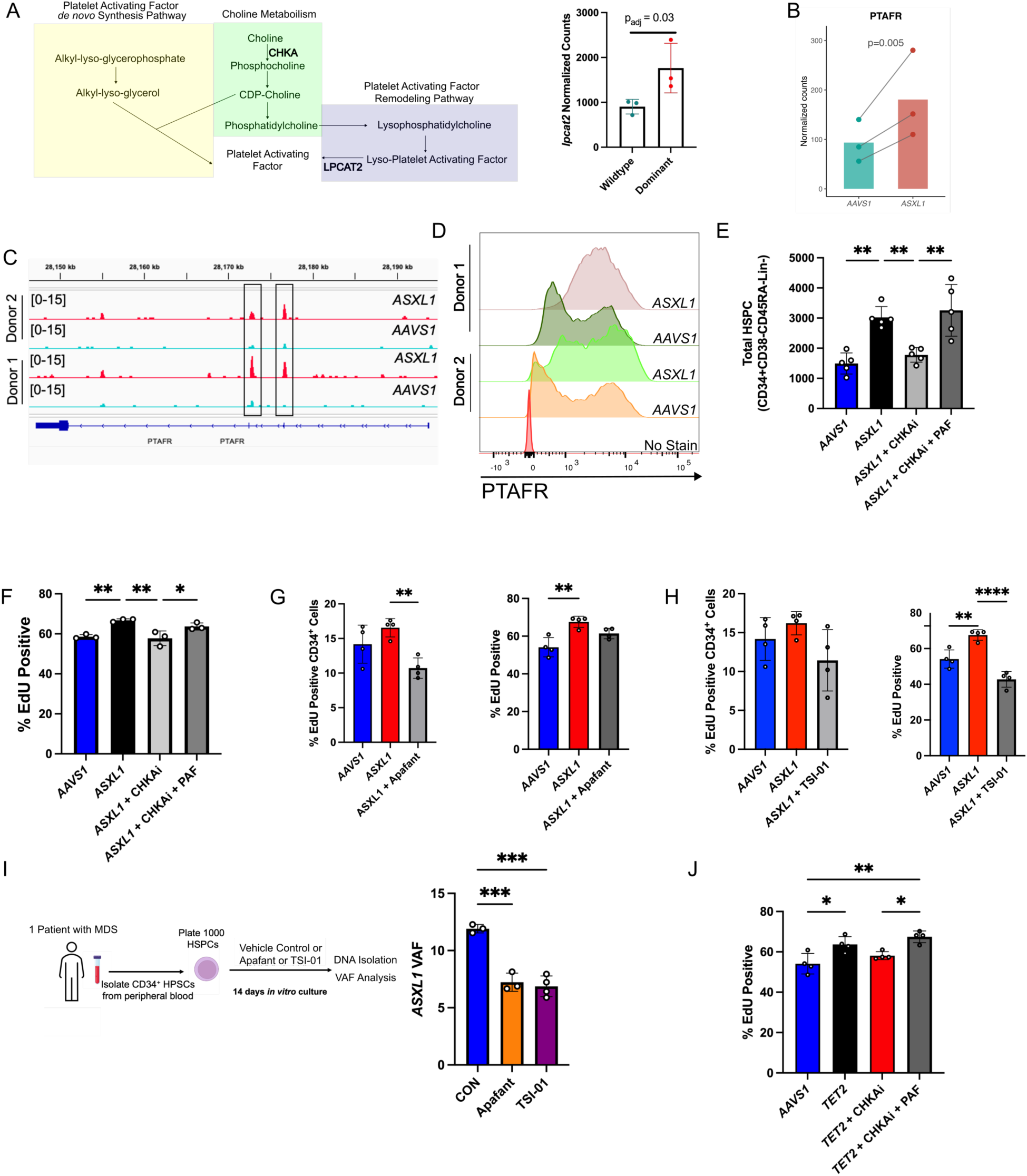
Demand for Platelet Activating Factor Underlies the Increased Phosphatidylcholine Synthesis in *ASXL1* and *TET2* Mutant HSPCs. A) Schematic outline of the intersection between choline and PAF metabolism with CHKA and LPCAT2’s relevant roles. Quantification of *lpcat2* gene expression differences between dominant, *asxl1*-mutant HSPC clones compared to non-dominant WT HSPC clones obtained from zebrafish. B) Quantification of *PTAFR* gene expression differences in primary, human HSPCs (CD34^+^) edited with CRISPR/Cas9 and a gRNA targeting a control region of the genome (*AAVS1*) or *ASXL1* collected 7 days after mutagenesis, n = 3 biological replicates. C) Gene tracks show regions of chromatin at *PTAFR* promoters that are open based on ATAC-seq in primary, human HSPCs (CD34^+^) edited with CRISPR/Cas9 and a gRNA targeting a control region of the genome (*AAVS1*) or *ASXL1* collected 7 days after mutagenesis, n = 2 biological replicates. D) Representative flow cytometry traces of PTAFR protein in primary, human HSPCs (CD34^+^) edited with CRISPR/Cas9 and a gRNA targeting a control region of the genome (*AAVS1*) or *ASXL1* collected 7 days after mutagenesis, n = 2 biological replicates. E) Total number of primary, human HSPCs (Lin^−^CD34^+^CD38^−^CD45RA^−^) 14 days after sorting 1000 cells per well in response to 1 μM CHKAi or 1 μM CHKAi and 100 nM PAF n = 5 technical replicates, **P < 0.01 (one-way ANOVA followed by Tukey’s Post Hoc). F) EdU positivity of cultures of HSPCs 4 days after *AAVS1* or *ASXL1* mutagenesis, EdU was incubated for 24 hours from day 3 to day 4, in response to 1 μM CHKAi and 100 nM PAF, n = 3 technical replicates representative of 3 independent experiments, *P < 0.05, **P < 0.01, (one-way ANOVA followed by Tukey’s Post Hoc). G) EdU positivity of cultures of HSPCs 4 days after *AAVS1* or *ASXL1* mutagenesis, EdU was incubated for 24 hours from day 3 to day 4, in response to 500 nM apafant, n = 4 technical replicates representative of 3 independent experiments, **P < 0.01, (one-way ANOVA followed by Tukey’s Post Hoc). H) EdU positivity of cultures of HSPCs 4 days after *AAVS1* or *ASXL1* mutagenesis, EdU was incubated for 24 hours from day 3 to day 4, in response to 10 μM TSI-01, n = 4 technical replicates representative of 3 independent experiments, **P < 0.01, ****P < 0.0001 (one-way ANOVA followed by Tukey’s Post Hoc). I) Experimental schematic of how human HSPCs were obtained from peripheral blood draws of a patient with myelodysplastic syndrome (MDS) and clonal dynamics evaluated in response to apafant or TSI-01. Quantification of *ASXL1* VAF change between DMSO control and apafant and TSI-01 14 days after treatment initiation, n = 3-4 technical replicates, ***P < 0.001 (one-way ANOVA followed by Tukey’s Post Hoc). J) EdU positivity of cultures of HSPCs 4 days after *AAVS1* or *TET2* mutagenesis, EdU was incubated for 24 hours from day 3 to day 4, in response to 1 μM CHKAi and 100 nM PAF, n = 3 technical replicates representative of 3 independent experiments, *P < 0.05, **P < 0.01, (one-way ANOVA followed by Tukey’s Post Hoc).

If the production of PAF by *ASXL1* mutant HSPCs required CHKA-dependent choline flux, then addition of exogenous PAF should be sufficient to overcome CHKAi induced clonal expansion reduction. Indeed, PAF was able to increase the total number of HSPCs (**Figure 4E**) and proliferation, as noted by EdU positivity (**Figure 4F**), 14 days and 4 days after *ASXL1* mutagenesis respectively. Modulation of PTAFR activity can be achieved through multiple modalities independent of influencing PC biosynthesis directly such as directly antagonizing PTAFR via the small molecule Apafant^34^ or through inhibiting LPCAT2 and subsequently the remodeling pathway with TSI-01^35^. Dose response experiments showed that proliferation of HSPCs from healthy donors was unaffected at the doses used for subsequent experiments (**Supplementary Figure 11AB**). Apafant treatment reduced proliferation of *ASXL1* mutant CD34^+^ HSPCs with no effect noted on the bulk cell population (**Figure 4G**) whereas TSI-01 was more effective at reducing the entire population of proliferating cells (**Figure 4H**). We obtained CD34^+^ from the peripheral blood of a patient with MDS in a manner similar to that described in **Figure 2H** and found that both Apafant and TSI-01 were sufficient to reduce *ASXL1* VAF after 14 days of culture (**Figure 4I**). Our findings demonstrate that PC/PAF axis modulation is effective in suppressing clonal expansion of cells from patients where the burden of disease has progressed to clinical diagnosis.

We next questioned whether this need for PAF signaling was also a feature of *DNMT3A* and *TET2* mutant HSPCs. We performed secondary analysis on an RNA-sequencing dataset of murine hematopoietic stem cells (Lin^−^Sca-1^+^c-Kit^+^CD48^−^CD150^+^) from different *Dnmt3a* genotypes including WT (*Dnmt3a*^+/+^), heterozygote (*Dnmt3a*^+/-^), knockout (*Dnmt3a*^−/-^), or two catalytically deficient mutants (*Dnmt3a*Δ/R832A, *Dnmt3a*Δ/E752A) and identified that knockout (p=0.06) and both catalytically deficient mutants had increased expression of *CHKA* (**Supplementary Figure 12A**). These mutants did not have increased expression of *Lpcat2* and *Ptafr* and therefore we did not pursue mechanistic studies related to this genotype (**Supplementary Figure 12B**). We propose that while all three of the most prominent CH-driving genotypes require enhanced CHKA activity PAF metabolism is not transcriptionally regulated in *DNMT3A* mutant cells like *ASXL1*.

We reanalyzed a previously published RNA-seq dataset where Lin^−^Sca-1^+^c-Kit^+^ (LSK) HSCs were isolated from mice harboring a *Tet2* catalytic-deficient allele (*Tet2*m), *Tet2* knockout (*Tet2*KO), or WT controls to determine levels of *Ptafr* and *Lpcat2*^36^. *Ptafr* expression increased (logFC = 1.68) in the LSK fraction in the *Tet2*KO cells specifically (**Supplementary Figure 12C**). *Lpcat2* was also increased in *Tet2*KO LSKs (logFC = 1.82) implicating the conserved effect of CH-mutations in enhancing the PC/PAF metabolism axis in *TET2* mouse models (**Supplementary Figure 12C**). Additionally, when bone marrow cells from *Tet2*KO or WT littermates controls were immortalized to a granulocyte-monocyte progenitor like fate (Lin^−^Sca-1^−^c-Kit^+^CD34^+^CD16/32^+^) through overexpression of *HOXB4 Lpcat2* was dramatically increased (**Supplementary Figure 12D**)^37^. Functionally, CHKAi abrogated the *TET2*-mutation driven proliferative advantage of human HSPCs which was rescued by PAF (**Figure 4J**) in a manner similar to *ASXL1* mutant human HSPCs (**Figure 2F**). Therefore, enhanced CHKA-dependent choline flux represents the first genotype conserved and small molecule responsive metabolic aberration in CH.

## Discussion

Our findings establish CHKA-dependent choline flux as a mutation agnostic metabolic dependency of the major genetic subtypes of CH. We combined color barcoding with low input metabolite and lipid profiling assays to identify a reduced abundance of betaine and acetylcholine in favor of choline flux through CHKA toward CDP-choline and subsequent PCs. We used stable isotope tracing studies combined with *ASXL1* mutagenesis in human HSPCs to show that enhanced CHKA activity is conserved in humans. We next showed that inhibition of CHKA suppressed clonal expansion across models of CH driven by *ASXL1*, *TET2*, and *DNMT3A*. These findings extend the concept of metabolic vulnerabilities in CH beyond individual genotypes and identify PC synthesis as a potential point of metabolic convergence among genetically diverse CH-driving clones.

A perplexity of CH has been how mutation in genes with diverse and at times opposing molecular functions (e.g., *DNMT3A* and *TET2*) converge on similar clonal expansion phenotypes. Our work serves to identify how genetic lesions in *ASXL1*, *DNMT3A*, and *TET2* all converge on a shared metabolic requirement. Importantly, this convergence does not appear to reflect identical downstream metabolic programs. It appears their downstream requirement of CHKA-dependent choline flux differs with *ASXL1* and *TET2* having an additional need to support PAF production. In contrast, *DNMT3A*-mutant HSPCs exhibit CHKA dependence without a corresponding increase in *LPCAT2* or *PTAFR* expression. A recent analysis found a positive correlation between CH status and circulating levels of Lp-PLA2, an enzyme that can degrade extracellular PAF, in *TET2* and *ASXL1* driven CH but not, *DNMT3A* driven CH^4^. This implies that while modulation of CHKA-dependent choline flux is a promising cross-genotype therapeutic target future studies should evaluate the utility of PAF signaling modulation in *DNMT3A* models.

Our findings extend observations from cancer metabolism to pre-malignant hematopoiesis. Enhanced choline metabolism is a hallmark of many types of cancer notably solid tumors^38–40^ to the extent that choline PET was approved for diagnostic purposes in recurrent prostate cancer^41,42^. Alongside the increase in choline metabolism in solid tumors is the repeated finding of the transcriptional upregulation of *CHKA*^43^. To exploit this choline dependency a series of molecules were developed to target CHKA with RSM-932a reaching phase I clinical trials^44,45^. These reports suggest the increased demand for PC is to satisfy the structural demands of rapid proliferation of cancer cells and necessity for lipids to make new cell membranes. In the hematopoietic system specifically, the choline transporters SLC44A1 and SLC44A2 are expressed to greater degrees in acute myeloid leukemia and acute lymphocytic leukemia cells compared to CD34^+^ cells from healthy donors^46^. This increase in choline demand was shown to facilitate choline to betaine catabolism which was suspected to support one-carbon metabolism and methylation reactions. Similarly, we show choline to betaine metabolism is enriched in *asxl1* mutant HSPCs however, we note one-carbon units are not liberated from betaine and therefore do not contribute to one-carbon metabolism in our model. Our findings recapitulate previous findings that betaine does not contribute to one-carbon metabolism in hematopoietic cells^47,48^. Our findings diverge from the previous literature in that we do not observe consistent transcriptional upregulation of *CHKA*. Increased dependence on PC synthesis does not necessarily require increased CHKA expression and may instead reflect altered pathway usage or increased metabolic demand.

We developed a patient-derived *ex vivo* clonal competition assay by isolating CD34^+^ HSPCs from peripheral blood of patients with clones harboring CH-driving mutations who presented with clonal cytopenia of undetermined significance (CCUS) or MDS. Because intra-patient clones are evaluated together, this approach preserves their competitive interactions. Using this model, we demonstrate the ability of CHKAi, Apafant, and TSI-01 to prevent clonal expansion of *ASXL1* mutant cells from people with advanced hematopoietic clonal disorders. By circumventing the need for mobilization or invasive marrow sampling this approach will offer an accessible method for testing interventions in patient derived samples at scale. The ability to derive these cultures directly from patients also provides a tractable approach for studying MDS, where genetic and clonal heterogeneity has complicated the development of experimentally tractable models and effective targeted therapies^49^. This new system provides a novel platform to test multiple therapeutics in a manner that faithfully recapitulates human disease.

The identification of CHKAi as a way to suppress expansion of disease-causing clones is exciting however, because CH affects such a large fraction of the population treatment of all individuals identified is not feasible. Future studies should center on identifying the population that is most likely to receive benefits from CHKAi. The ability to further stratify risk of individuals with CH such as the Clonal Hematopoiesis Risk Score^50^ or MN-predict^51^ will aid in the identification of people who may most benefit from CHKAi as a therapeutic intervention. Altogether, our data reveal a metabolic architecture in which genetically diverse CH-driving HSPC clones converge on increased CHKA-dependent choline flux while still retaining genotype-specific downstream phenotypes.

## Methods

### Zebrafish models

Wildtype Tübingen (TU), *draculin:CreERT2* (referred to as *drl:CreERT2*)^52^, and *Ubi:Zebrabow-M* (referred to as *ubi:Zebrabow*)^18,53^ were used in this study. All animals were housed at the Boston Children’s Hospital with all experiments being performed according to protocols approved by the Institutional Animal Care and Use Committee (IACUC, Protocol Number: 00002043).

### Mouse Models

This study uses C57BL/6 (Black 6) Fgd5*^ZsGr-CreERT^*^227^, Dnmt3a*^fl-R^*^878^*^H^*^26^, and Rosa26*^lox-STOP-lox-tdTomato^* mice^54^. Both Dnmt3a^fl-878H^ and Fgd5^ZsGr-CreERT2^ alleles are maintained as heterozygotes in experimental mice. For *in vitro* culture studies long term hematopoietic stem cells (Lin-Sca-1^+^cKit^+^CD48^−^CD150^+^) cells were isolated by flow cytometry. Antibodies for analysis were sourced as follows: DAPI (for dead cell exclusion; Sigma-Aldrich); CD3, CD11b, Ter119, CD19, B220, Gr1, cKit (Invitrogen); Sca1, CD150, CD127 (Biolegend); CD48 (BD Biosciences). Lineage (Lin) cocktail includes CD3, CD11b, CD19, B220, Ter119, and Gr1. Isolated Long-term hematopoietic stem cells were cultured as previously described in a polyvinyl alcohol-based media^28^. Cells were seeded at 1×10^3 cells per well and periodically passaged (every 2-3 days) by light mechanical dissociation without chemical or enzymatic aids for re-plating in fresh fibronectin coated plates (Corning). After 5 days, cells were treated with 500 nM 4-OHT (Sigma-Aldrich) to induce recombination and expression of tdTomato and *Dnmt3a^R^*^878^*^H^*.

For *in vivo* transplantation experiments, LT-HSCs were isolated from Fgd5*^CreER^* Rosa26*^lox-STOP-loxtdTomato^* Dnmt3a*^fl-R^*^878^*^H^* mice as described above. After 7 days in culture, cells were treated with 500 nM 4-OHT and tdTomato expression was quantified via microscopy. Images were captured on a Carl Zeiss inverted microscope or Nikon 90i. For each well, at each timepoint, 4 random fields were imaged in brightfield, and dsRed channels. ImageJ was used to convert images and export each channel as a separate image for quantification. Cell segmentation and quantification in brightfield and fluorescent channel images was performed in Cellpose using the built-in Cyto model for cell counting^55^. The tdTomato+ value for each well is an average across all four fields collected. 2.5×10^5 cells were transplanted into C57BL/6J mice that were lethally irradiated with two 500cGy doses separated by 3 hours. All mice were 3-4 months of age at transplant. Experiments were performed according to approved protocols by the Institutional Animal Care and Use Committee at the Boston Children’s Hospital (IACUC, Protocol Number: 00002284).

### Mobilized Peripheral Blood CD34+ HSPCs

Human CD34+ HSPCs from G-CSF mobilized peripheral blood of healthy volunteers were procured from the Fred Hutchinson Cancer Research Center. HSPCs were maintained in StemSpan II media (STEMCELL Technologies), with 10X CD34+ Expansion supplement (STEMCELL Technologies) and 0.1% Penn/strep (Gibco) at 37oC and 5% CO2.

### Non-Mobilized Peripheral Blood CD34+ HSPCs

5-10 mL of peripheral blood was collected in EDTA tubes through the Adult Acute Leukemia Program at the Dana Farber Cancer Institute, with Institutional Review Board approval and informed patient consent in accordance with the Declaration of Helsinki (IRB Number: 01-206 for collection and biobanking and 25-786 for secondary use clinical data). Patients included had a diagnosis of CCUS or MDS and available tissue. Clinical data were anonymized and samples were de-identified. Mononuclear cells were washed twice with PBS (Fisher) and 2% FBS (R&D Systems) followed by centrifugation at 300 x g for 8 minutes at room temperature. Cells were passed through a 35 μm nylon mesh strainer (Fisher) followed by centrifugation at 300 x g for 10 minutes at room temperature and aspiration of the supernatant. The pellet was resuspended in 150 μL MACS buffer (PBS, 0.5% BSA, 2mM EDTA), 50 μL FcR Blocking Reagent and 50 μL CD34 Microbeads (Miltenyi Biotec) and incubated at 4°C for 30 minutes. Cells were centrifuged at 300 x g for 10 minutes at room temperature and resuspended in 500 μL MACS buffer. LS Columns (Miltenyi Biotech) were placed on a MACS Separator and prepared by rinsing with 3 mL MACS buffer. Cells were passed through the column with unlabeled flow-through collected for DNA isolation. Three washes of 3 mL MACS buffer were performed. The LS column was removed from the separator and the CD34^+^ HSPCs were flushed with 1 mL MACS buffer using the provided plunger. Isolated cells were cultured in HSPC Expansion Media (StemSpan II media (STEMCELL Technologies), with 10X CD34+ Expansion media (STEMCELL Technologies) and 0.1% Penicillin/Streptomycin (Gibco) at 37°C and 5% CO_2_.

### Tissue editing With Inducible Stem cell Tagging via Recombination (TWISTR)

The TWISTR system was employed as previously described^17^. *drl:CreERT2* and *ubi:Zebrabow* lines were crossed, clutches divided and embryos injected with purified gRNAs to drive CH or controls and Cas9 mRNA as described below. Initial TWISTR experiments included injections of pools of gRNAs targeting genes including *dnmt8*, *dnmt6*, *asxl1*, *tet2*, *ezh2*, *stag2a*, *stag2b*, *rad21a*, *tp53*, *zrsr2, piga* and/or control genes (*albino* or *golden*). For subsequent experiments *asxl1* was injected with *albino* or *albino* was injected alone to generate WT controls. Color recombination was induced 24 hours post fertilization (hpf). 25-35 embryos were transferred to a single well of a 6-well plate and treated with 15 μM 4-hydroxytamoxifen (4-OHT, Sigma-Aldrich) for 4 hours protected from light at 28.5°C before being returned to fresh embryo buffer.

### CRISPR-Cas9 of CD34+ human HSPCs

CD34+ HSPCs were prescreened for *TCL1A* rs2887399 using ThermoFisher SNP assay (Assay ID: C 15842295_20) to ensure they were WT. After thawing, HSPCs were allowed to recover in the HSPC Expansion media described above for 48 hours prior to gene editing. 3.2 μg of gRNA of *AAVS1*, *TET2*, or *ASXL1* (Synthego) and 8.18 μg Alt-R S.p. Cas9 Nuclease V3 (IDT) were incubated for 15 min at room temperature with light mixing to form a ribonucleoprotein complex prior to electroporation. Guide sequences and associated primers can be found in **Supplementary Table 4**. HSPCs were resuspended in 18 μL of P3 solution (Lonza) and mixed with 2 μL of gRNA/Cas9 ribonucleoprotein complex and transferred to 16-well Nucleocuvette strips (Lonza) and electroporated using a Lonza 4D Nucleofector (program DZ-100). Cells were allowed to recover in the Nucleocuvette for 10 minutes prior to being transferred to 1 mL of HSPC expansion media for 24 hours. After 24 hours DNA isolation was performed using a DNA blood and tissue kit (Qiagen) and editing efficiency measured using sanger sequencing and ICE Analysis^56^.

### Zebrafish Flow Cytometry Analysis and Isolation of Dominant Clones

Zebrafish were sacrificed in ice-cold fish water and marrows isolated under a Leica MZ75 light microscope. Kidney marrows were placed in PBS (Gibco) with 2% FBS and 1 USP units/mL heparin (Sigma-Aldrich). To create a single-cell suspension kidney marrows were mechanically dissociated via repeated pipetting and passed through a 35 μm nylon mesh strainer. Flow cytometry lineage analysis was performed on a BD LSRII or Fortessa (BD Biosciences) and cell type (myelomonocytes, HSPCs, lymphocytes, and erythrocytes) were defined based on their forward and side light scatter characteristics as previously defined^57^. DRAQ7 was used at a concentration of 3 nM for viability analysis (Abcam). For Zebrabow color analysis a BD FACSAria II with special order 445 nm laser for CFP detection was used (BD Biosciences). Data was analyzed using FlowJo^TM^ v10 (BD Biosciences) followed by the previously published Zbow software^17,58^. Only fish with a red only clone of less than 25% indicating 75% recombination were retained for analysis. Clones were initially identified using the myelomonocytes gate to best represent recent output.

### Human CD34+ HSPC Liquid Expansion and Pharmacological Treatment

For total HSPC counting assays cells were prepared for FACS 24 hours after gene editing. Cells were stained with an extracellular HSPC marker panel: CD34 (APC-CD34, Clone 561, 1:50 dilution, BioLegend), CD38 (PE/Cy7-CD38, Clone HIT2, 1:100 dilution, BioLegend), CD45RA (BV605-CD45RA, Clone HI100, 1:40 dilution, BioLegend), and a lineage cocktail (PE/Cy5-CD2, Clone RPA-2.10, 1:100 dilution, BioLegend; PE/Cy5-CD3, Clone HIT3a, 1:100 dilution, BioLegend; PE/Cy5-CD4, Clone SK3, 1:100 dilution, BioLegend; PE/Cy5-CD8a, Clone HIT8a, 1:100 dilution, BioLegend; PE/Cy5-CD16, Clone 3G8, 1:100 dilution, BioLegend; PE/Cy5-CD19, Clone HIB19, 1:100 dilution, BioLegend, PE/Cy5-CD20, Clone 2H7, 1:100 dilution, BioLegend; PE/Cy5-CD56, Clone 5.1H11, 1:100 dilution, BioLegend; PE/Cy5-CD235a, Clone HIR2, 1:400 dilution, BioLegend; PE/Cy5-CD14, Clone M5E2, 1:100 dilution, BioLegend) in 100 μL of PBS, 2% FBS, and 1 mM EDTA. 500-1000 CD34^+^Lin^−^CD38^−^CD45RA^−^(HSPCs) were sorted into 100 μL HSPC Expansion Media into a single well of a 96-well plate using a BD FACSAria II and Fixable Viability Stain 700 was used to eliminate dead cells (BD Biosciences). Four days after sorting 100 μL HSPC Expansion Media was added, 10 days after sorting cells were transferred to a 48-well plate along with 400 μL additional HSPC Expansion Media. Fourteen days after sorting cells were stained with the HSPC marker panel as above with counting beads and total HSPCs were quantified using a BD FACSAria II followed by analysis in FlowJo^TM^ v10 (BD Biosciences). For EdU experiments cells were plated at 25 000 cells per 400 μL HSPC Expansion Media in a single well of 48-well plate directly after gene editing. For analysis taking place 7 days after gene editing (e.g., metabolite profiling, lipid profiling, RNA-seq, ATAC-seq) cells were plated at a density of 100 000 in 1 mL HSPC Expansion Media in a well of 24-well plate and transferred with an additional 3 mL HSPC Expansion Media to 1 well of a 6-well plate after 4 days of culture.

Drug and lipid treatments were initiated 24 hours after gene editing or sorting to allow cells to recover. All compounds were purchased from commercially available vendors (choline chloride trimethyl-D_9_, Cambridge Isotope Laboratories, Inc,; RSM-932a, MedChemExpress; propargylcholine, Cayman Chemicals; V-11-0711, MedChemExpress; platelet activating factor C16, Tocris; Apafant, MedChemExpress; TSI-01, Cayman Chemicals) used at the doses specified in their respective figures. Whenever new media was added, fresh chemical was added to the new media to retain the same overall drug concentration.

### *In vivo* Stable Isotope Tracing

To measure active choline metabolism 5-10 zebrafish per liter were submerged in fish water containing 1 mM d9-choline off system flow. To maintain oxygenation while the tank was off system flow air was continuously pumped into the tank. Fish water containing fresh d9-choline was provided after 24 hours.

### Zebrafish Intraperitoneal Injections

Adult zebrafish (6-24 months post fertilization, mpf) were anesthetized using 0.02% tricaine and injected using a 26-gauge Hamilton syringe delivering the doses of chemicals detailed in the figures or their vehicle controls both diluted in PBS. Intraperitoneal injection was performed in the abdominal cavity, posterior to the pelvic girdle Syringes were cleaned between washes with 70% ethanol and H_2_O. Zebrafish were allowed to recover in fresh fish water before being returned to their tanks.

### Zebrafish Retroorbital Bleeds

Zebrafish 6-24 mpf were anaesthetized in 0.02% tricaine in fish water. A 26-gauge Hamilton syringe was used to procure blood from the retroorbital sinus and placed in 200 μL PBS, 2% FBS and 1 USP units/mL heparin. DNA was extracted using the Blood and Tissue DNAeasy kit (Qiagen).

### Human HSPC Colony Forming Unit Assay

Twenty-four hours after electroporation cells were incubated with RSM-932a or vehicle control in HSPC Expansion Media. Two days after incubation with RSM-932a HSPCs were plated at a density of 1000 cells/mL in H4034 methylcellulose medium (STEMCELL Technologies) according to the manufacturer’s instructions. After 14 days of incubation at 37°C with 5% CO_2_ colony quantification was performed using STEMVision instrument.

### Non-Mobilized Peripheral Blood HSPC Competition Assay

Two days after magnetic bead isolation 1000 HSPCs were plated in 100 μL HSPC Expansion Media into a single cell of a 96-well plate and drug treatment initiated. Four days after sorting 100 μL HSPC Expansion Media was added, 10 days after sorting cells were transferred to a 48-well plate along with 400 μL additional HSPC Expansion Media. Fourteen days after drug initiation cells were counted using a hemocytometer after incubation in trypan blue dye. Cells were also collected for DNA isolation via a Blood and Tissue Kit DNA extraction kit (Qiagen).

Primers were made for each patient’s unique genetic mutations. A 10 μL PCR reaction containing 5 μL 2X Phusion High-Fidelity PCR Master Mix (NEB), 1.25 μL forward primer (10 μM), 1.25 μL reverse primer (10 μM), 2 μL DNA template, 1 μL nuclease-free H2O. For each amplicon one primer was synthesized with one of 14 custom barcode sequences appended to the 5’ end allowed for 14 reactions to be multiplexed. Cycling conditions were 98°C for 3 min; 35 cycles of 98°C for 10 sec, 60°C for 10 sec, 72°C for 10 sec, and a final extension at 72°C for 5 min.

Barcoded PCR amplicons were pooled and PCR purification was performed using a commercially available kit (Qiagen). Pooled samples were sequenced on the Illumina Miseq. Samples were analyzed using the CRISPResso2 package^59^. Variant allele frequency was determined by quantifying the number of reads including the mutant sequence divided by the total number of reads (i.e., mutant reads/[mutant reads + reference reads]).

### Variant Allele Frequency in Zebrafish

For variant allele frequency as a surrogate as clonal tracking in zebrafish DNA from retroorbital bleeds were used. The CRIPSR loci of *asxl1* was amplified using a 10 μL PCR reaction containing 5 μL 2X Phusion High-Fidelity PCR Master Mix (NEB), 1.25 μL forward primer (10 μM), 1.25 μL reverse primer (10 μM), 2 μL DNA template, 1 μL nuclease-free H2O. For each amplicon one primer was synthesized with one of 14 custom barcode sequences appended to the 5’ end allowed for 14 reactions to be multiplexed. Cycling conditions were 98°C for 3 min; 35 cycles of 98°C for 10 sec, 60oC for 10 sec, 72°C for 10 sec, and a final extension at 72oC for 5 min. After PCR purification the product was sequenced using the Illumina Miseq platform. Mutations were analyzed using a previously published protocol as described^17^.

### Zebrafish Phosphatidylcholine Profiling

To minimize lipid contamination a BDFACS Aria II was primed to replace all sheath fluid with PBS exclusively. Single-color HSPC clones were sorted, as described, into PBS containing 1% charcoal stripped FBS (ThermoFisher), centrifuged at 300 x g for 5 min, supernatant removed and snap frozen in liquid nitrogen. Cells were then lysed in a buffer to quench metabolism, as previously described^60^. The lysed cells were subjected to a Bligh-Dyer extraction. The nonpolar (chloroform) fraction was used to analyze intact lipids on a Thermo Fusion Lumos liquid chromatography-high resolution mass spectrometry (LC-HRMS) system. Analysis was performed using the TraceFinder 5.2 software platform from Thermo Scientific, as described^61^.

### Human HSPC Phosphatidylcholine Profiling

CD34+ HSPCs were isolated 7 days after mutagenesis using the magnetic purification method described above. From day 5-7 cells were incubate with d9-choline. Cells were then lysed in a buffer to quench metabolism, as previously described^60^. The lysed cells were subjected to a Bligh-Dyer extraction. The nonpolar (chloroform) fraction was used to analyze intact lipids on a Thermo Fusion Lumos liquid chromatography-high resolution mass spectrometry (LC-HRMS) system. Analysis was performed using the TraceFinder 5.2 software platform from Thermo Scientific, as described^61^ and D9-integration calculated.

### Human HSPC Polar Choline Metabolite Profiling

CD34+ HSPCs were isolated 7 days after mutagenesis using the magnetic purification method described above. From day 5-7 cells were incubate with d9-choline. Cells were then lysed in a buffer to quench metabolism, as previously described^60^. The Bligh-Dyer extraction was used to obtain aqueous. The aqueous (methanol) fraction was analyzed on an Exploris liquid chromatograph – high resolution Orbitrap mass spectrometer to detect choline related metabolites.

### Whole Genome Bisulfite Sequencing of Dominant Zebrafish HSPCS

Dominant and WT HSPC clones (50 000 total cells per replicate) were isolated as described and DNA was isolated using the PureLink^TM^ Genomic DNA mini Kit (Invitrogen). Whole genome bisulfite sequencing was performed as previously described^62,63^. Differentially methylated regions were identified using the Metilene pipeline^64^ using a predefined set of parameters including: 1) distance between CpGs within the same DMR were less than 300 base pairs, 2) a minimum of 10 CpGs per DMR was required, and 3) dinucleotides had greater than 10 reads per sample.

### EdU Proliferation Analysis

Proliferation via EdU incorporation was measured using the Click-iT^TM^ Plus EdU Alexa Fluor^TM^ 488 Flow Cytometry Assay Kit (ThermoFisher) according to the manufacturer’s protocol. 10 μM EdU was added to human HSPC culture for 24 hours prior to harvest. For experiments including concurrent CD34 staining cells were incubated with CD34+ prior to fixation and downstream processing for the Click-iT reaction.

### Phosphatidylcholine Synthesis Assay

A click chemistry assay to measure phosphatidylcholine synthesis was developed based on previously described methods^65^. Cells were incubated with 4 mg/L propargylcholine (Cayman Chemical) for 24 hours to enable metabolic incorporation of the alkyne-tagged choline analog. After the 24 hours of incubation cells were pelleted by centrifugation (300 x g for 5 min) in a FACS buffer (PBS, 2% FBS, and 0.5 mM EDTA). Cells were then fixed in 100 μL 4% PFA for 15 min at room temperature, protected from light. Importantly, cells were not permeabilized so that the click reaction occurred on the cell membrane giving a direct readout of membrane bound choline derived lipids and not other, intracellular polar metabolites. Cells were then washed in FACS buffer. The click reaction was performed in accordance with the manufacturer’s instructions described in the Click-iT^TM^ Plus Alexa Fluor^TM^ Picolyl Azide Toolkit (ThermoFisher Scientific).

### Platelet Activating Factor Protein Staining

Cells were washed in FACS buffer at the time points denoted in the figure legends, and fixed in 100 μL 4% paraformaldehyde for 15 min at room temperature and protected from light. Cells were then permeabilized in 1 mL ice cold 100% methanol for 30 min. After another wash in FACS buffer, cells were resuspended in 100 μL FACS buffer containing 2 μL of a FITC-conjugated PTAFR antibody (ABIN7347983) for 30 min at 4°C protected from light. After a wash in FACS buffer cells were analyzed on a FACS LSRII.

### Low-input Bulk RNA-sequencing of Dominant Zebrafish HSPCs

Dominant and WT HSPC clones (10 000 total cells per replicate) were isolated as described and RNA was obtained using the RNAeasy Plus Micro Kit (Qiagen). Ribosomal RNA was depleted from using the RiboGone-Mammalian kit (Takara), and cDNA libraries prepared using the SMARTer Universal Low Input RNA kit (Takara). cDNA quality was assessed using a TapeStation high-sensitivity D1000 ScreenTape assay (Agilent). Libraries were prepared using the ThruPLEX DNA-seq Kit (Takara). Libraries were sequenced on an Illumina HiSeq 4000. Raw reads were quality controlled with FastQC, and adapter sequences and low-quality regions were trimmed using Cutadapt. High-quality reads were aligned to the zebrafish genome (GRCz11) using the STAR 2.7.0 using Spliced Transcripts Alignment tool and gene-level raw read counts were quantified with HTSeq^66,67^.

### Bulk RNA-sequencing of Human HSPCS

After CD34+ magnetic bead enrichment RNA was isolated using a Direct-zol RNA Miniprep kit (Zymo). Purified RNA was shipped on dry ice to Plasmidsaurus. Library preparation and sequencing was performed by Plasmidsaurus using their 3’-end counting RNA-seq service via the Illumina platform.

Fastq file quality was assessed using FastQC v0.12.1 and reads were quality filtered using fastp v0.24.0 with poly-X tail trimming, 3’ quality-based tail trimming, a minimum Phred quality score of 15, and a minimum length requirement of 50 bp. Quality filtered reads were aligned to the human genome (GRCh38.p14) using STAR aligner v2.7.11 and coordinate sorting was performed using samtools v.1.22.1. UMIcollapse v1.1.0 was used to remove PCR and optical duplicates. Alignment quality metrics, read distribution, and strand specificity across genomic features were assessed using RSeQC v5.0.4 as well as Qualimap v2.3. Differential gene expression was done with edgeR V4.0.16.

### Assay for Transposase Accessible Chromatin (ATAC-seq)

50 000 human, CD34+ HSPCs were isolated by magnetic bead separation as described above. Cells were washed once in 50 μL ice cold PBS and re-pelleted at 500 x g for 5 min at 4°C. Cell pellets were lysed in 50 μL ATAC-seq lysis buffer (10 mM Tris-HCL pH 7.4, 10 mM NaCl, 3 mM MgCl_2_, 0.1% IGEPAL CA-630) followed by centrifugation at 500 x g for 10 min at 4°C. The transposition reaction was performed on ice in 25 μL 2X TD buffer, 2.5 μL Tn5 transposase (Illumina), and 22.5 μL nuclease-free H2O for 1 hour at 37oC. Transposed DNA was purified with the MinElute PCR Purification Kit (Qiagen) and eluted in 10 μL total volume.

Library construction followed the Illumina Nextera protocol using transposed DNA, NEB PCR master mix, SYBR Green, and universal and library-specific Nextera index primers. Initial PCR pre-amplification was performed (72°C for 5 min, 98°C for 30 sec, then 5 cycles of 98°C for 10 sec, 63°C for 40 sec, and 72°C for 1 min, hold at 4oC). Additional amplification cycle number was determined by a side qPCR on a 5 μL aliquot of the of the pre-amplified reaction (98°C for 30s, 20 cycles of 98°C for 10 secs, 63°C for 30 secs, and 72°C for 1 min, hold at 4°C). Individual libraries were then amplified for the empirically determined number for additional cycles (98°C for 30 sec, *n* cycles of 98°C for 10 sec, 63°C for 30 sec, and 72°C for 1 min, hold at 4°C). Final libraries were assessed for quality on a TapeStation high-sensitivity D1000 ScreenTape assay (Agilent) followed by sequencing on an Illumina HiSeq 2500.

For analysis ATAC-seq reads were aligned to the human genome (GRCh38.p14) using Bowtie v2.2.1^68^ with parameters --end-to-end, -N0, -L20. Coverage tracks for visualization were generated using MACS with parameters -w -S --space=50 –nomodel --shiftsize=200. Peaks were called using MACS2 v.2.10^69^ with parameters --nomodel -- shift -100 --extsize 200 and a q-value enrichment threshold of 0.05 across all datasets.

### Secondary Analysis of Publicly Available RNA-seq Datasets

#### Dnmt3a

Raw gene count and FPKM values from GO accession GSE272913^70^. Murine hematopoietic stem cells (Lin^−^Sca-1^+^c-Kit^+^CD48^−^CD150^+^) from different *Dnmt3a* genotypes including WT (*Dnmt3a*^+/+^), heterozygote (*Dnmt3a*^+/-^), knockout (*Dnmt3a*^−/-^), or two catalytically deficient mutants (*Dnmt3a*Δ/R832A, *Dnmt3a*Δ/E752A) were compared. From the deposited count files per-sample FPKM values were loaded and for each gene of interest and FPKM ± standard error of the mean was computed per genotype. Comparisons between each mutant group and control were performed using a two-tailed t-test and log2 fold changes were calculated.

#### Tet2

Data was obtained from GEO accession GSE132090^71^ which contained bulk RNA-seq profiles of Lin^−^Sca-1^+^c-Kit^+^ (LSK) HSCs isolated from mice harboring a *Tet2* catalytic-deficient allele (*Tet2*m), *Tet2* knockout (*Tet2*KO), or WT controls. Differential expression analysis was performed using DESeq2. Wald test results were used and log2 fold change and Benjamin-Hochberg adjusted p-values were reported. For a second *Tet2* data set FPKM values were obtained from GEO accession GSE264041^72^. From the deposited count files per-sample FPKM values were loaded and for each gene of interest and FPKM ± standard error of the mean was computed per genotype.

Comparisons between each mutant group and control were performed using a two-tailed t-test and log2 fold changes were calculated.

### Statistical Analysis

Graphs and statistical analysis were produced using RStudio or Prism v10 GraphPad Software, Inc.). For all graphs the mean +/-standard deviation is displayed. Statistical tests include 1-way ANOVA, 2-way ANOVA, two-sided paired tests, and two-sided unpaired t-tests. The specific tests, p-values, and post hoc analysis are described in relevant figure legends.

## Acknowledgements

L.I.Z is supported by the HHMI Investigator Program. This work was supported by the National Institutes of Health P01HL131477 (L.I.Z.). B.J.G received support from the Canadian Institutes of Health Research (458577), The Prevent Cancer Foundation, and the National Institutes of Health (K99AG086600). Research reported in this publication was supported by the National Institute of Diabetes and Digestive and Kidney Diseases of the National Institutes of Health under Award Number U24DK126127. The content is solely the responsibility of the authors and does not necessarily represent the official views of the National Institutes of Health. Peripheral blood sample collection supported by the Ted and Eileen Pasquarello Tissue Bank in Hematological Malignancies. R.J.D. is supported by the HHMI Investigator Program, N.C.I. grant R35CA220449, and the Eugene McDermott Distinguished Professorship for the Study of Human Growth and Development. The Children’s Research Metabolomics Core Facility was supported by CPRIT grant RP180778.

---

**Supplementary Figure 1.**
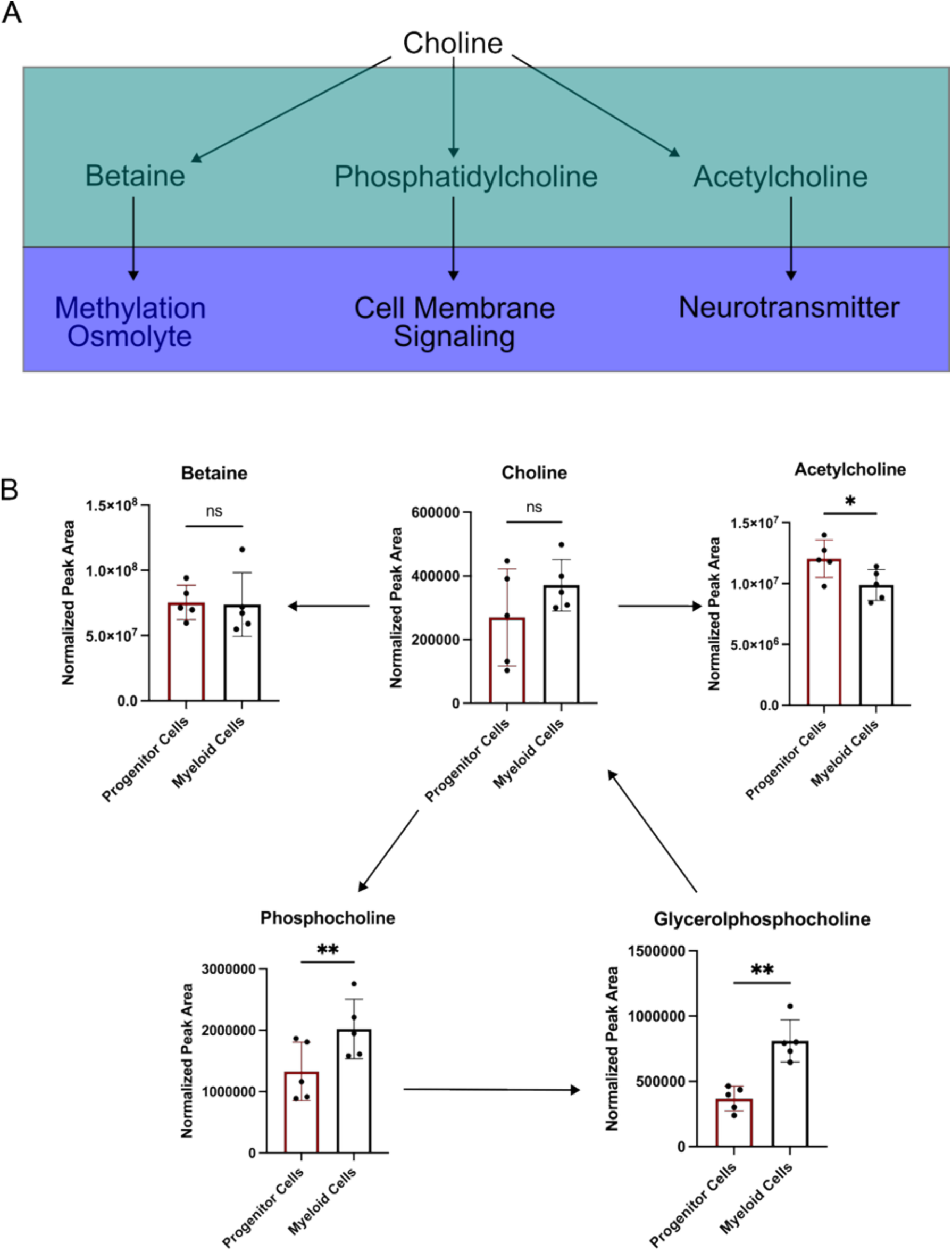
Quantification of Choline Metabolism in Hematopoietic Zebrafish Cells. A) Schematic of the intracellular fates of choline and their downstream function. B) Levels of choline and choline-derived metabolites in zebrafish HSPCs and myelomonocytes. n = 5 biological replicates, *P < 0.05, **P < 0.01 (unpaired t-test).

**Supplementary Figure 2.**
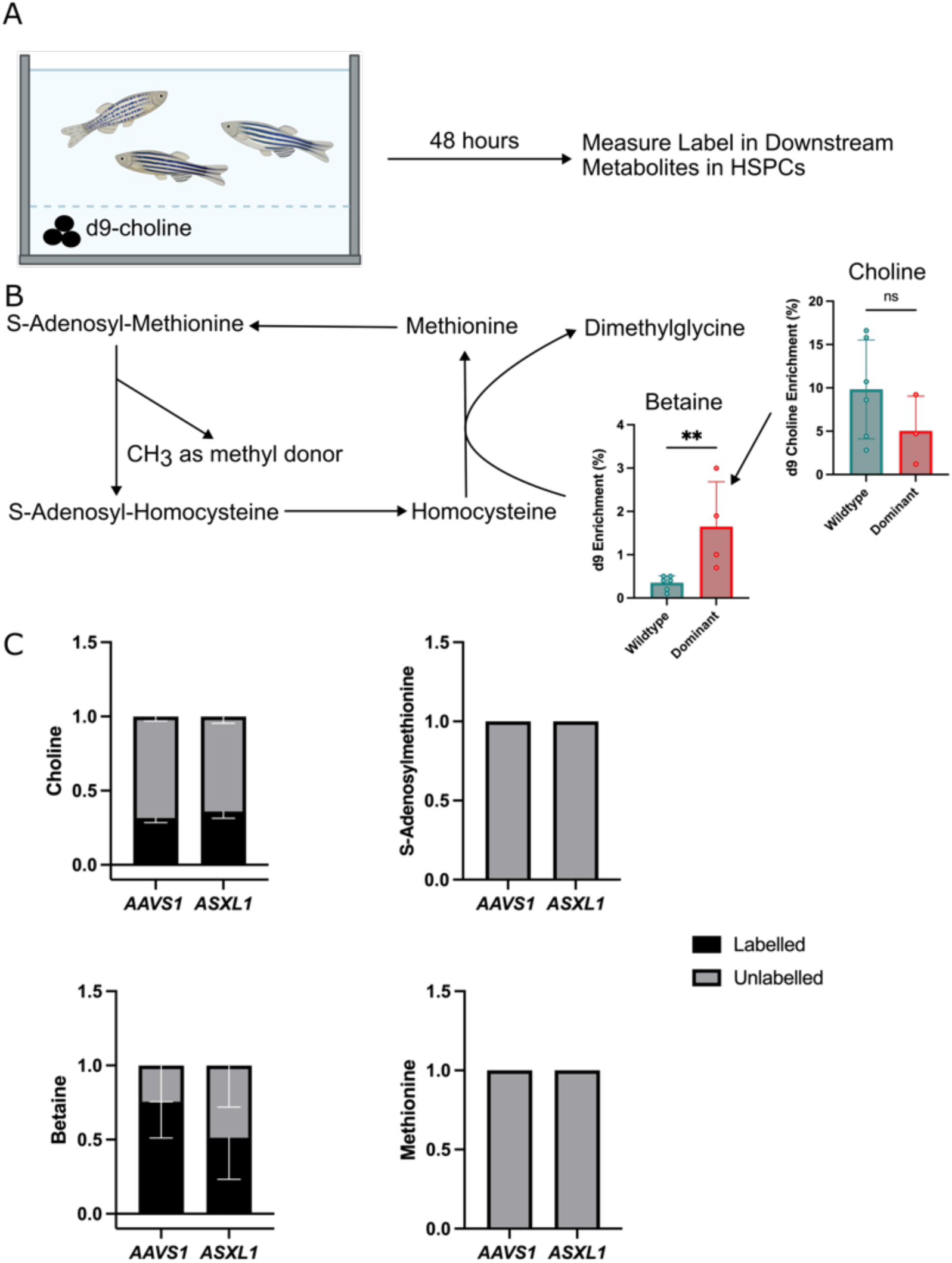
Betaine Biosynthesis is Altered in Dominant, *asxl1-*Mutant HSPCs. A) Schematic of *in vivo* d9-choline stable isotope tracing approach. B) Schematic of how choline-betaine catabolism can support one-carbon metabolism. Dot plots show enrichment of deuterated (labelled) choline in choline and betaine. n = 4-6 biological replicates, **P < 0.01 (unpaired t-test). C) Stacked bar graphs represent fraction of each one-carbon related metabolite labelled by deuterated choline in primary, human HSPCs edited with CRISPR/Cas9 and a gRNA targeting a *AAVS1* or *ASXL1*. n = 4 biological replicates, (paired t-test).

**Supplementary Figure 3.**
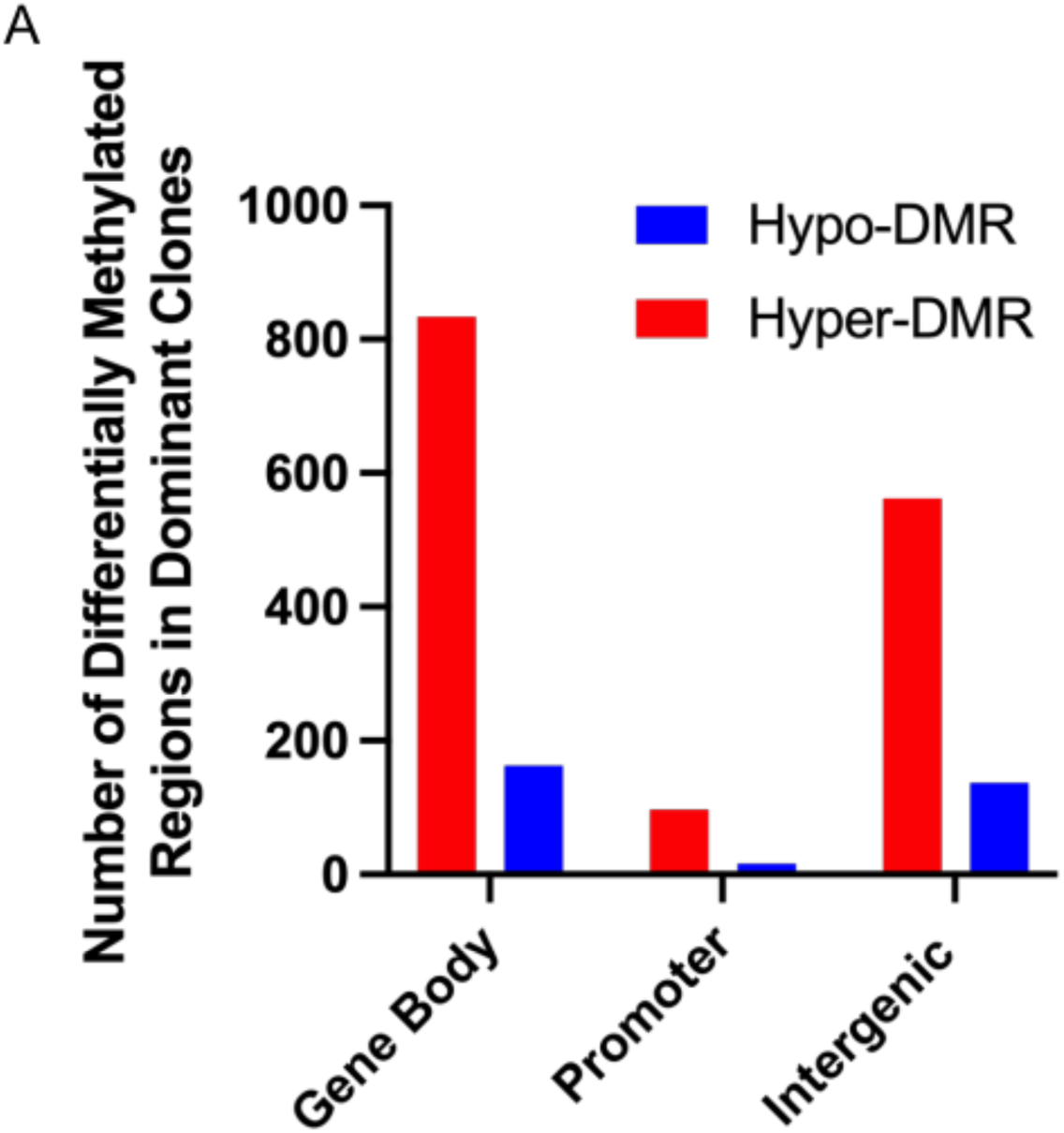
DNA Methylation Profile of Dominant, *asxl1-mutant* HSPCs. A) Abundance of differentially methylated regions (DMRs) in dominant, *asxl1*-mutant HSPCs compared to WT controls in the gene body, promoter, and intergenic. n = 3 biological replicates.

**Supplementary Figure 4.**
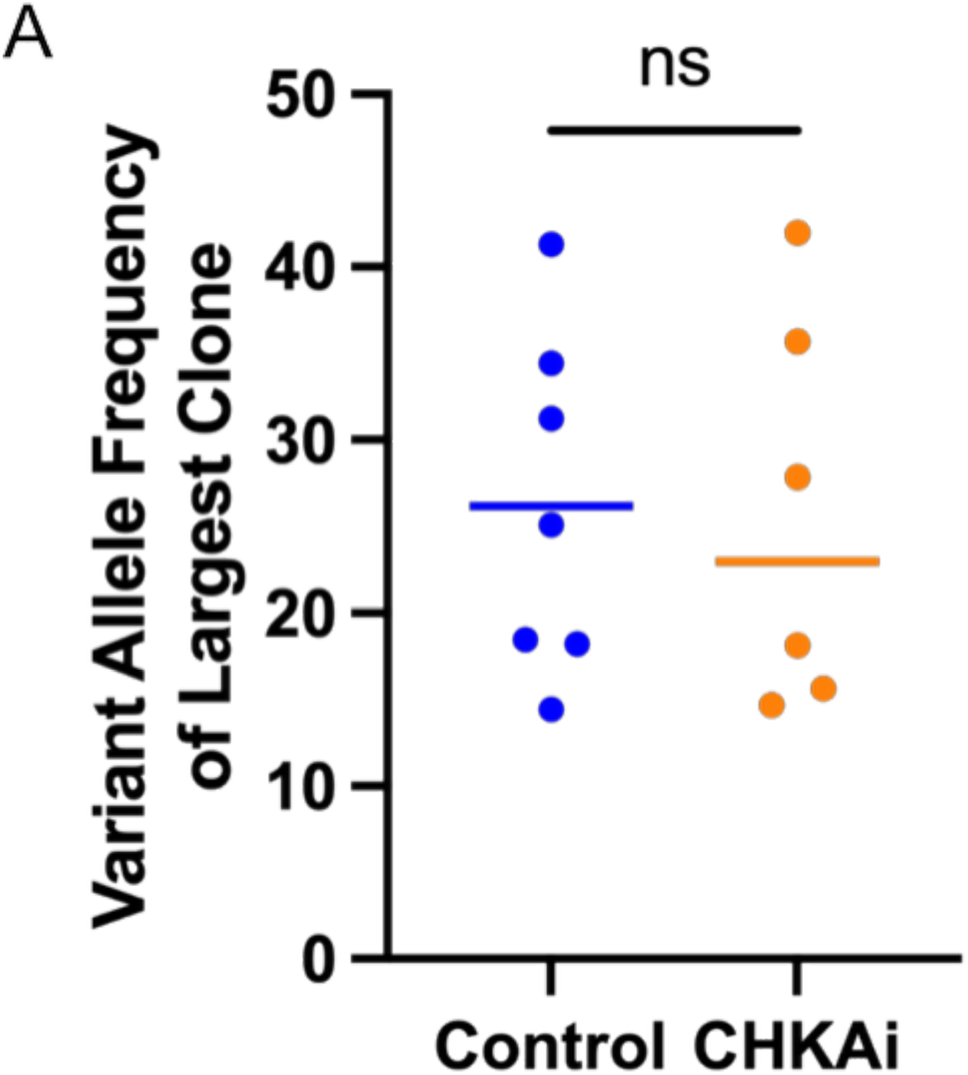
Size of the Largest *asxl1-*Mutant Clone Prior to CHKAi Treatment in Zebrafish. A) Quantification of the size of the largest *asxl1* variant allele frequency (VAF) in the peripheral blood prior to initiation of treatment of PBS control or CHKAi, n = 6-7 biological replicates, (unpaired t-test).

**Supplementary Figure 5.**
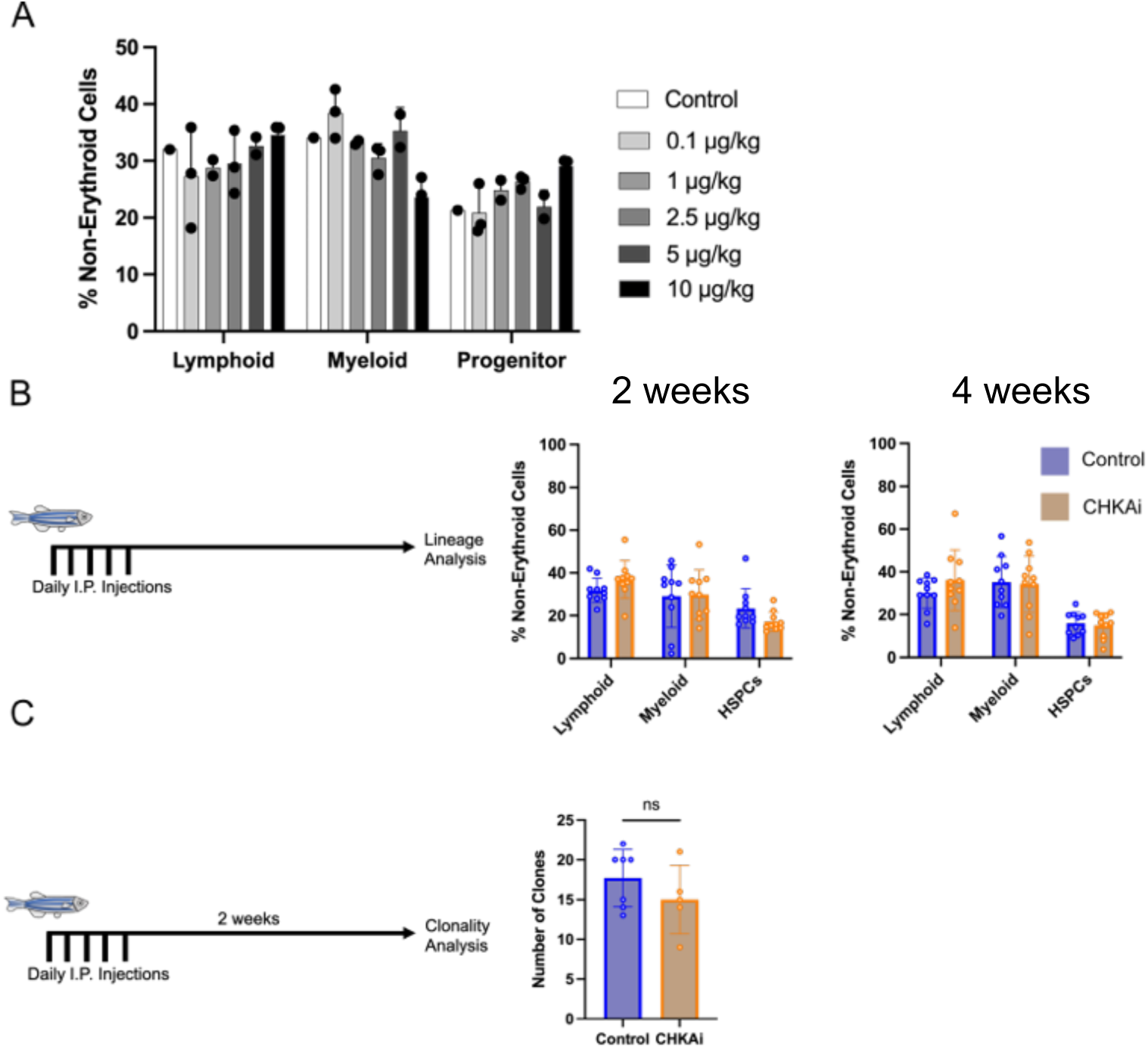
Characterization of CHKAi Treatment on Hematopoiesis in a Wildtype Setting. A) Quantification of hematopoietic lineage output based on flow cytometry of forward and side scatter light characteristics in wildtype zebrafish 1 week after exposure to a range of CHKAi doses (0-10 μg/kg), n = 1-3 biological replicates. B) Quantification of hematopoietic lineage output based on flow cytometry of forward and side scatter light characteristics in wildtype zebrafish 2 and 4 weeks after 10 μg/kg CHKAi treatment, n = 10 biological replicates, (unpaired t-test). C) Number of clones contributing to hematopoiesis as measured by unique color barcodes in the myelomonocyte compartment of Zebrabow zebrafish 2 weeks after treatment with 10 μg/kg CHKAi, n = 5-7 biological replicates, (unpaired t-test).

**Supplementary Figure 6.**
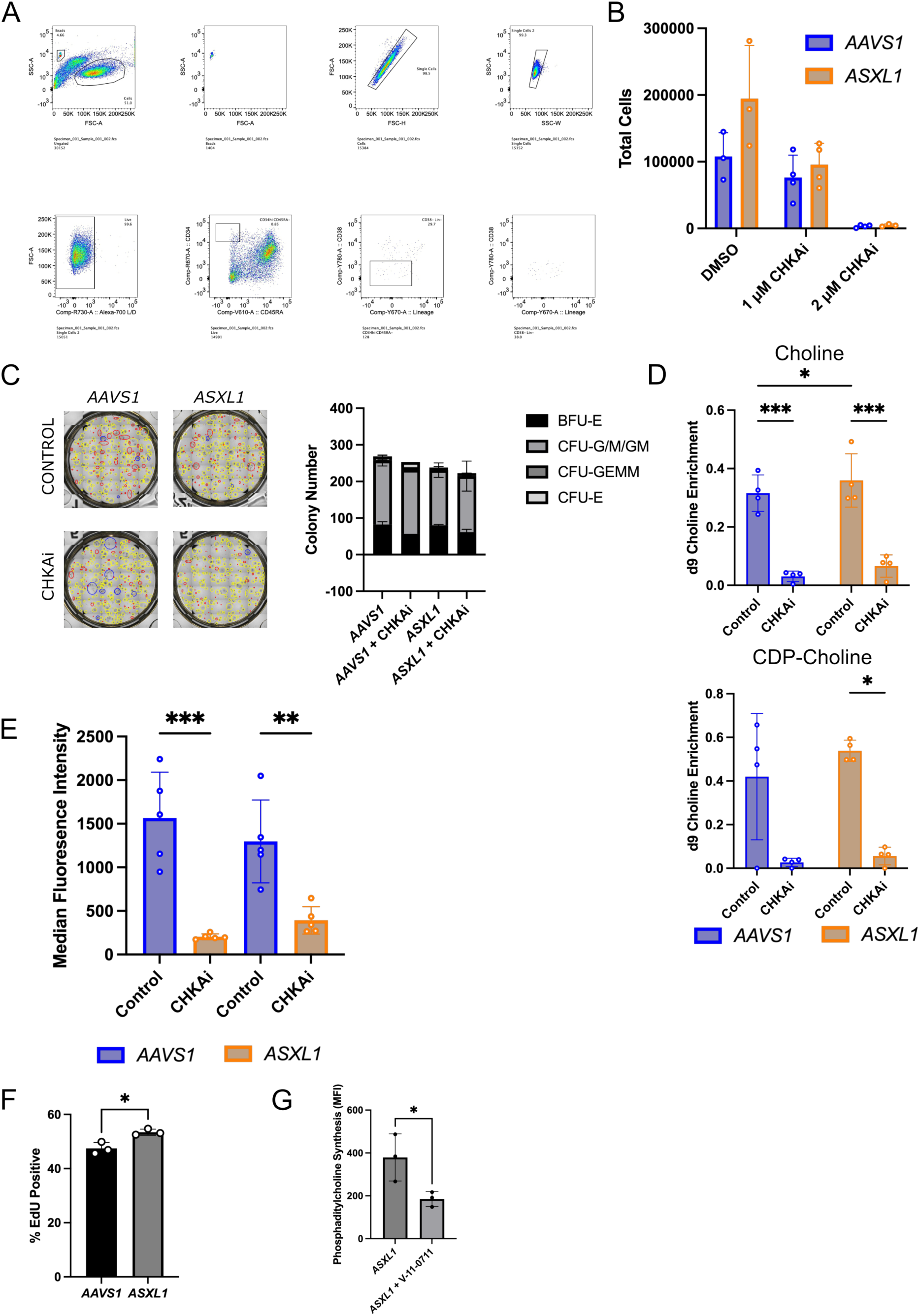
Choline Kinase Alpha Inhibition Has No Negative Effects on Wildtype Hematopoietic Cells. A) Representative fluorescence activated cell sorting strategy for isolating and quantifying HPSCs (Lin^−^CD34^+^CD38^−^CD45RA^−^). B) Quantification of total cells after 14 days exposure to escalating doses of CHKAi, n = 3-4 technical replicates. C) Representative image and quantification of colony number and type 1000 primary, human HSPCs (*AAVS1* or *ASXL1* mutant) were plated in MethoCult semisolid methylcellulose with or without 1 μM CHKAi, n = 3 technical replicates. D) Levels of integration of d9-choline into choline and CDP-choline in primary, human HSPCs (CD34+ enriched) after 7 days of culture in drug treatment, n = 4 biological replicates, *P < 0.05, ***P < 0.001. (two-way repeated measures ANOVA followed by Tukey’s Post Hoc). E) Quantification of membrane phosphatidylcholine synthesis as determined by click labeling after 48 hours of incubation with an alkynyl derivative of choline with or without CHKAi, n = 5 technical replicates, **P < 0.01, ***P < 0.001. (two-way repeated measures ANOVA followed by Tukey’s Post Hoc). F) EdU positivity of cultures of HSPCs 4 days after *AAVS1* or *ASXL1* mutagenesis, EdU was incubated for 24 hours from day 3 to day 4 n = 3 technical replicates representative of 3 independent experiments, *P < 0.05. (unpaired t-test). G) Quantification of membrane phosphatidylcholine synthesis as determined by click labeling after 48 hours of incubation with an alkynyl derivative of choline with or without 1 uM V-11-0711 in *ASXL1*-mutant human HSPCs, n = 3 technical replicates, *P < 0.05. (unpaired t-test).

**Supplementary Figure 7.**
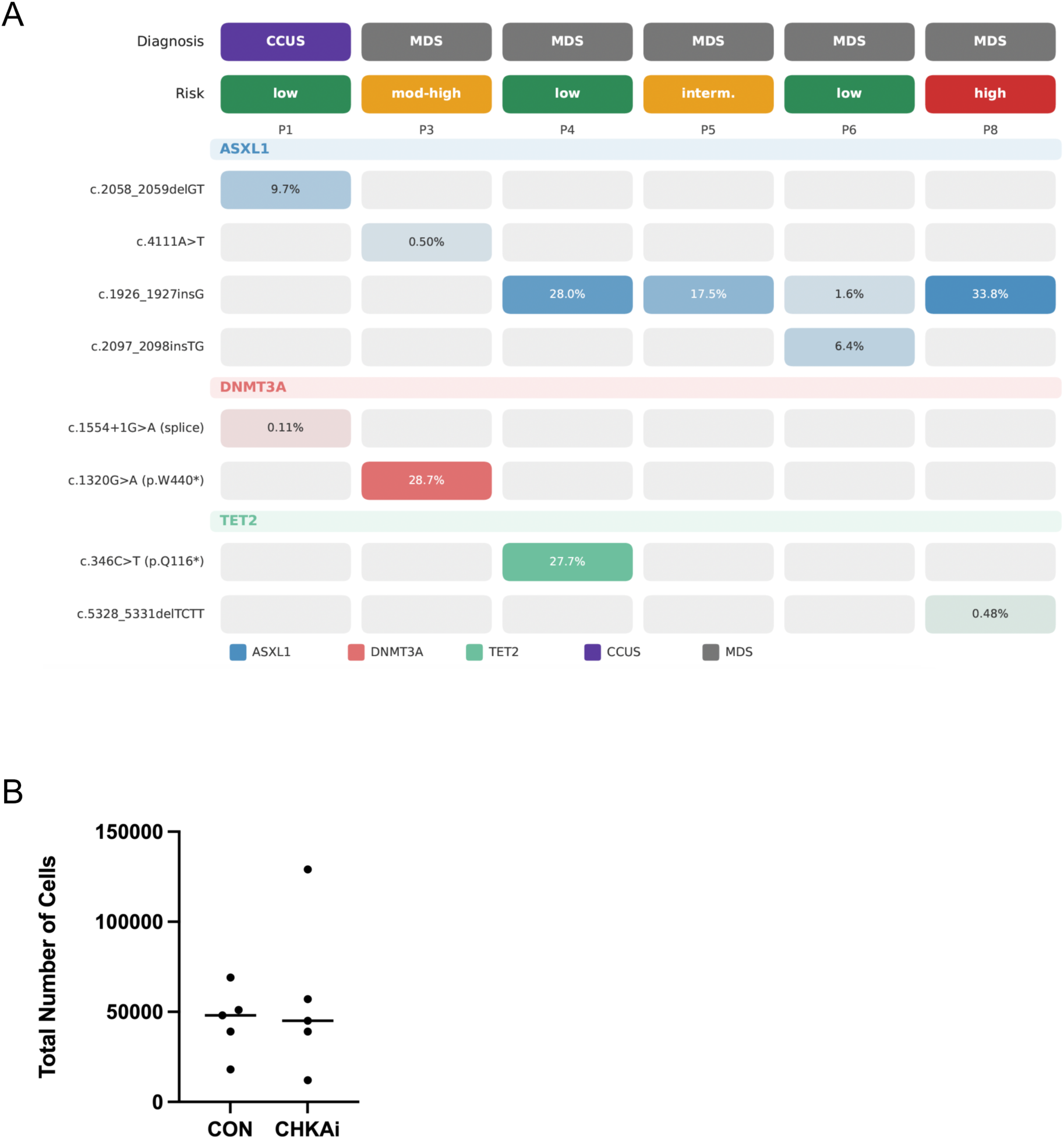
Characterization of Primary, Human HSPCs from Non-Mobilized Peripheral Blood. A) Oncoprint representation of clinical characteristics and variant allele frequencies of *ASXL1*, *DNMT3A*, *TET2* in patients from which CD34^+^ HSPCs were obtained in Figure 2H. B) Quantification of total cell number of HSPCs isolated from the peripheral blood of a healthy donor without mobilization after 14 days of culture with control or 1 μM CHKAi, n = 5 technical replicates. (unpaired t-test).

**Supplementary Figure 8.**
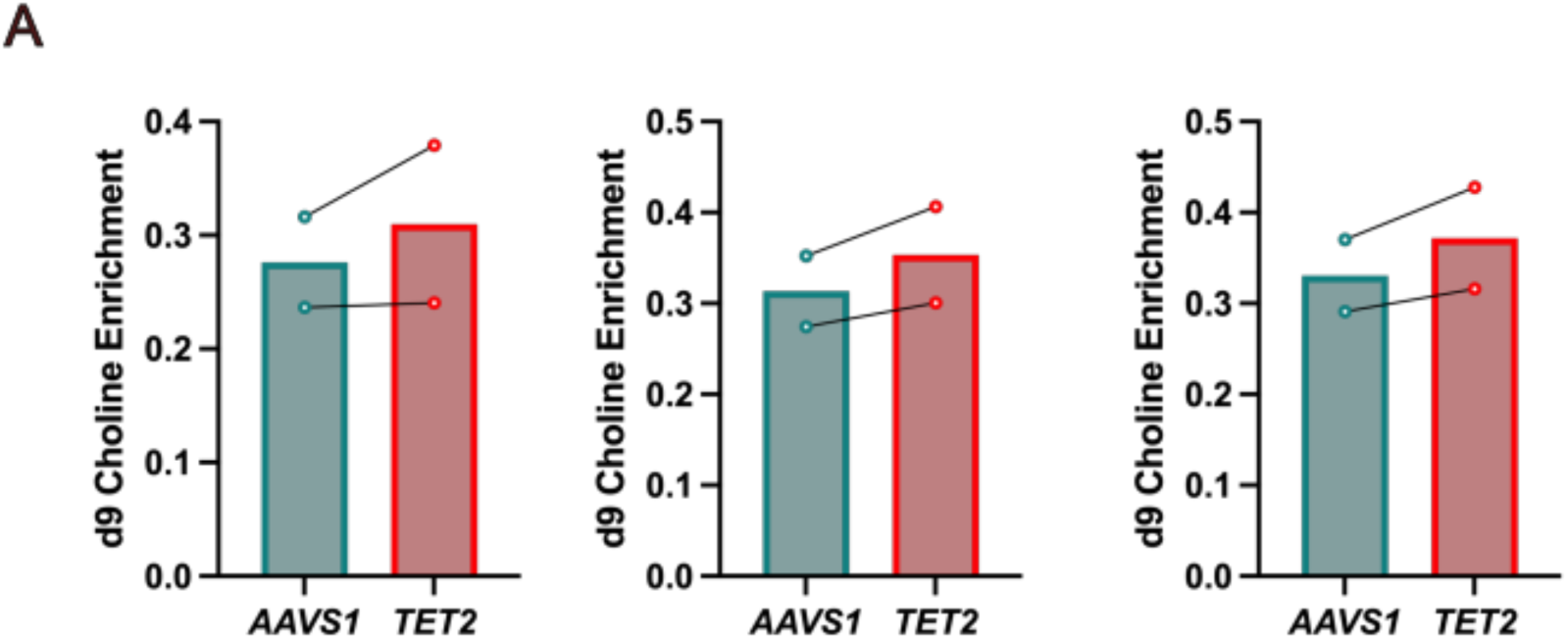
Effect of *TET2* Mutagenesis in Primary, Human HSPCs on PC Biosynthesis. A) Levels of integration of d9-choline into phosphatidylcholine species 16:0/16:0, 16:0/18:2, 16:1/18:1 in primary, human HSPCs edited with CRISPR/Cas9 and a gRNA targeting a control region of the genome (*AAVS1*) or *TET2*, n = 2 biological replicates.

**Supplementary Figure 9.**
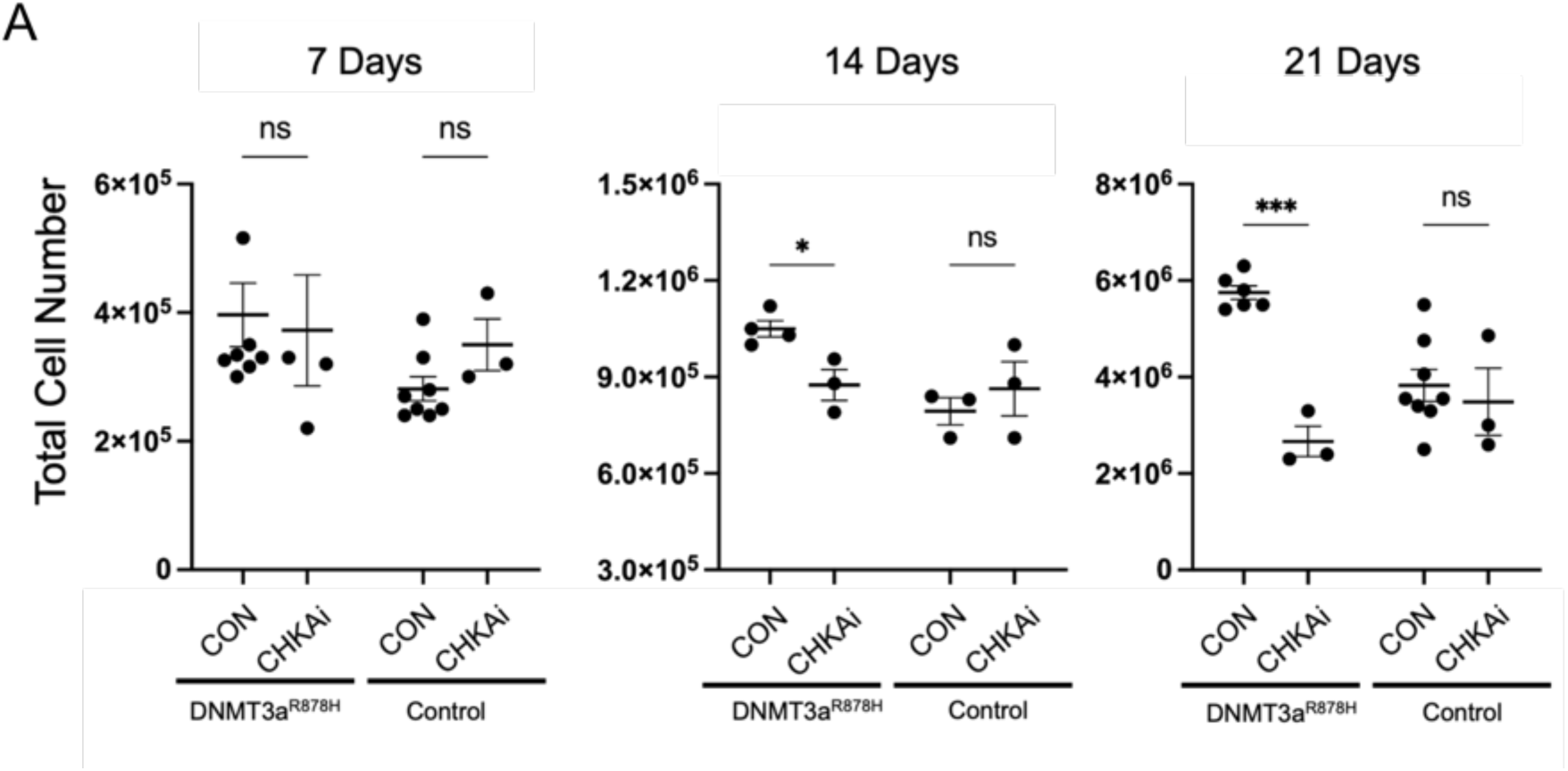
Choline Kinase Alpha Inhibition Reduces Proliferation in Dnmt3a^R878H/+^ Cells Specifically. A) Quantification of total cell number from cultures of HSPCs (Lin-Sca-1+cKit+CD48-CD150+) expanded *ex vivo* after induction of Dnmt3a^878H/+^ and tdTomato expression in isolated mice HSPCs (Lin-Sca-1+cKit+CD48-CD150+) in response to 1 μM CHKAi, n = 3-7 technical replicates, *P < 0.05, *** P< 0.01 (unpaired t-test).

**Supplementary Figure 10.**
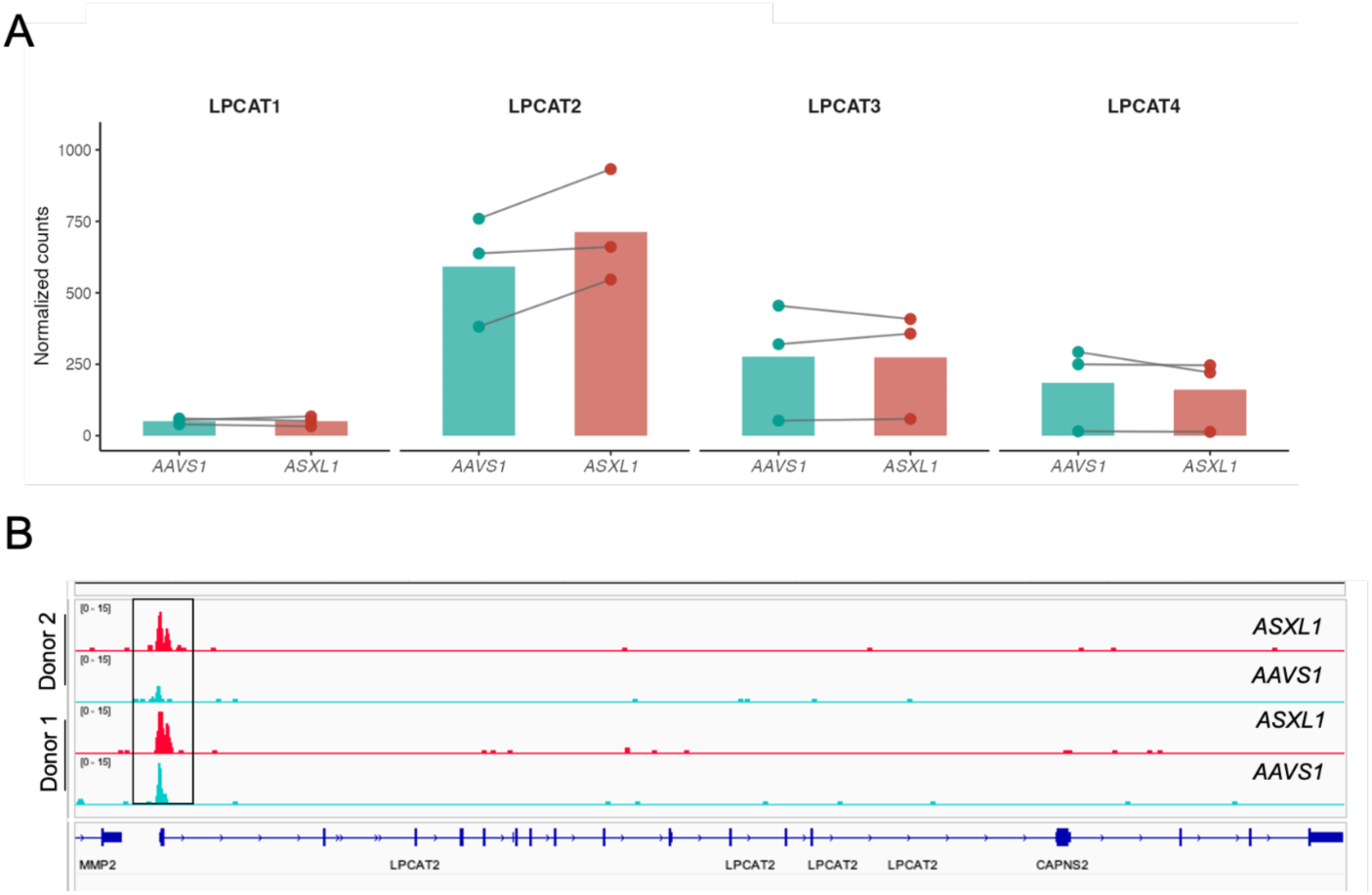
Profile of LPCAT2 Expression in primary, human *ASXL1*-mutant HSPCs. A) Quantification of *LPCAT* family gene expression differences in primary, human HSPCs (CD34^+^) edited with CRISPR/Cas9 and a gRNA targeting a control region of the genome (*AAVS1*) or *ASXL1* collected 7 days after mutagenesis, n = 3 biological replicates. B) Gene tracks show regions of chromatin on *LPCAT2* based on ATAC-seq in primary, human HSPCs (CD34^+^) edited with CRISPR/Cas9 and a gRNA targeting a control region of the genome (*AAVS1*) or *ASXL1* collected 7 days after mutagenesis, n = 2 biological replicates.

**Supplementary Figure 11.**
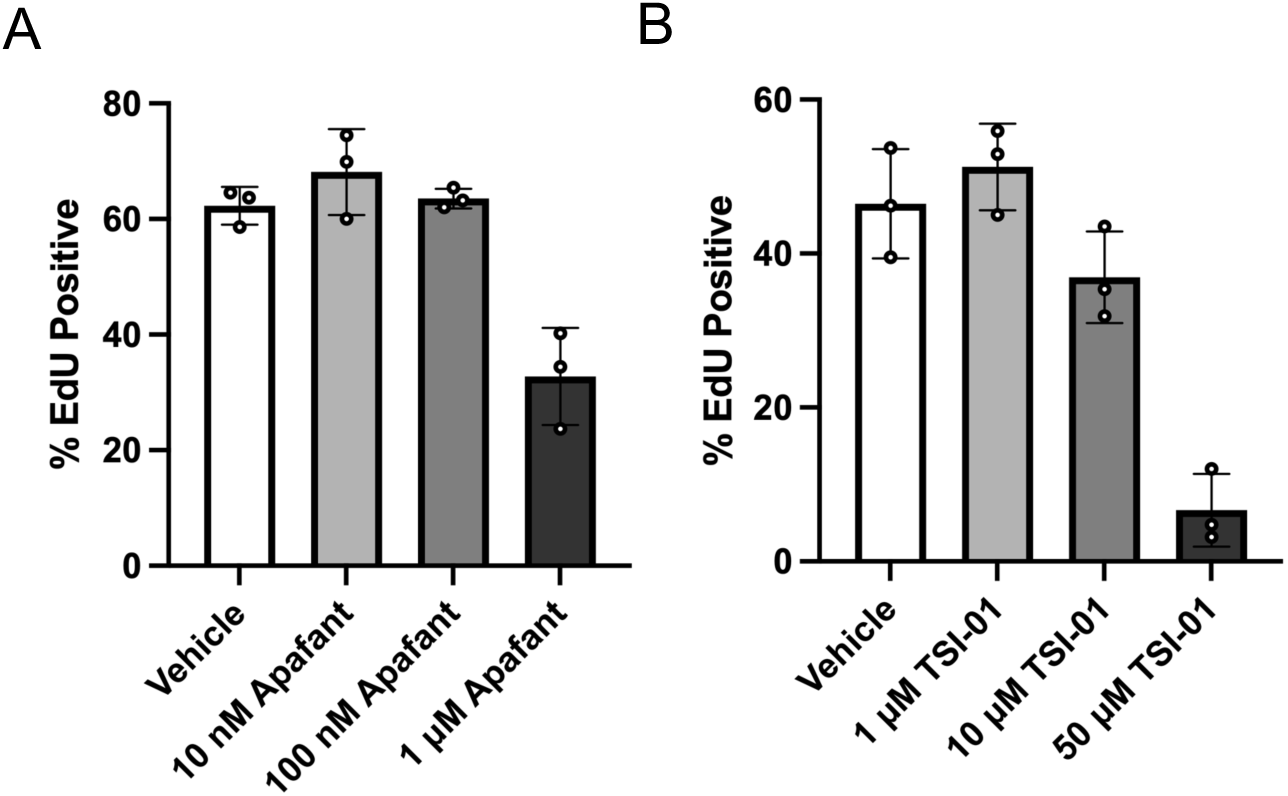
Dose Response of PAF Signaling Modulating Compounds. A) EdU positivity of cultures of HSPCs EdU was incubated for 24 hours from day 3 to day 4, in response to 10 nM, 100 nM, or 1 μM apafant, n = 3 technical replicates. B) EdU positivity of cultures of HSPCs EdU was incubated for 24 hours from day 3 to day 4, in response to 1, 10, or 50 μM TSI-01, n = 3 technical replicates

**Supplementary Figure 12.**
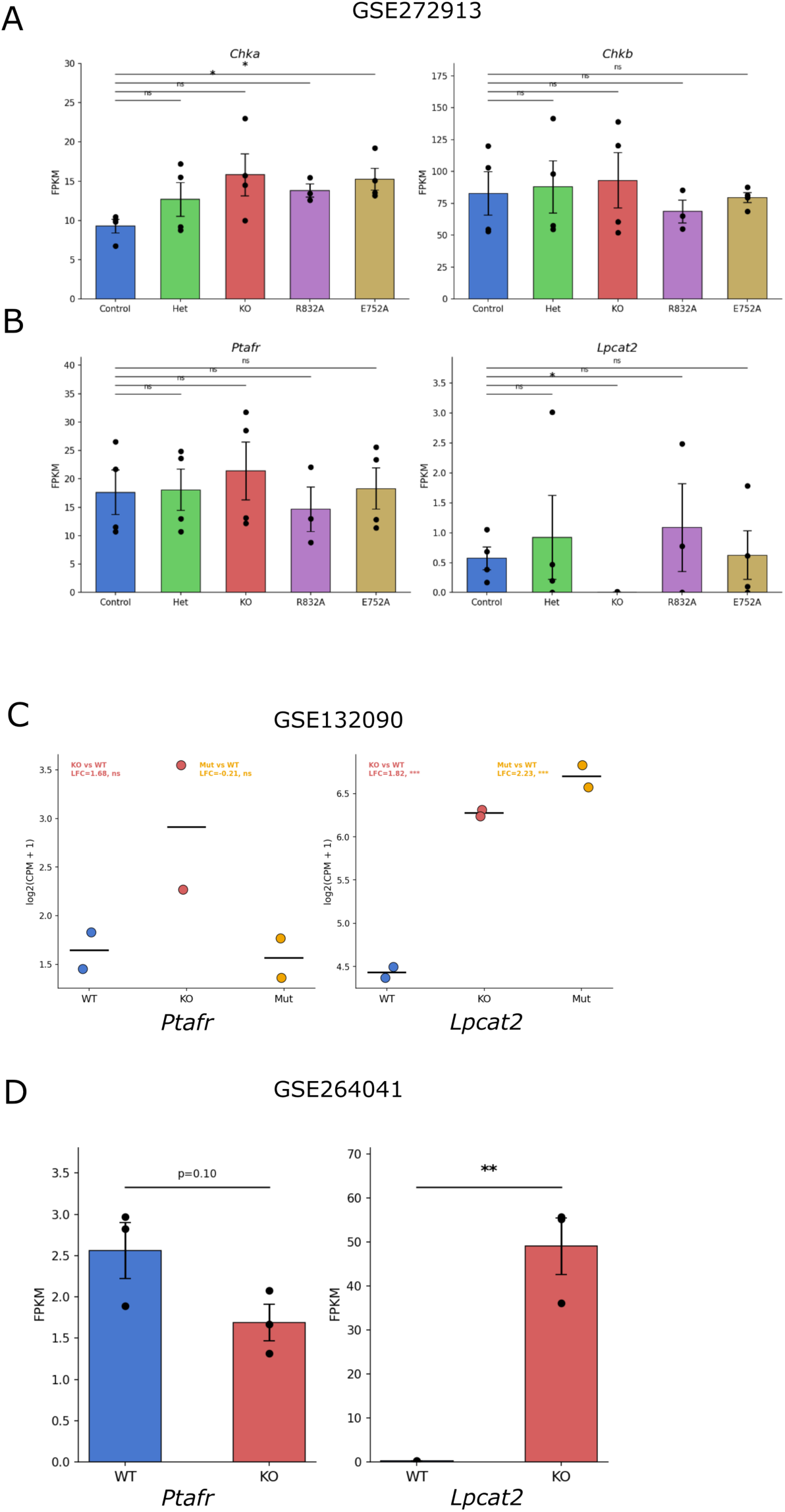
Transcriptional Profiling of PC and PAF Related Genes in *Tet2* and *Dnmt3a* mutant models. AB) Re-analysis of publicly available RNA-sequencing datasets for murine *Dnmt3a* loss of function HSPC models with *Chka, Chkab, Ptafr,* and *Lpcat2* quantified. CD) Re-analysis of publicly available RNA-sequencing datasets for murine *Tet2* loss of function HSPC models with *Ptafr* and *Lpcat2* quantified.

